# A Single-Cell Transcriptomic Atlas of Human Heart Organoids across Differentiation Protocols

**DOI:** 10.64898/2026.11.25.769101

**Authors:** Sarah Ouologuem, Vladana Đaković, Sophie Zengerle, Jannik Geissner, Tatjana Dorn, Anna B. Meier, Lika Drakhlis, Chrysoula Makra, Leonard Sima, Laura Xourgias, Julia L. Beckmann, Tristan Aretz, Arne van den Berg, Sina Mohrenweis, Korbinian Traeuble, Manuel Gander, Laura D. Martens, Karl-Ludwig Laugwitz, Robert Zweigerdt, Lennard Halle, Matthias Heinig, Alessandra Moretti, Fabian J. Theis

## Abstract

Human pluripotent stem cell-derived heart organoids have emerged as powerful models of human cardiac development, disease, and therapeutic responses. Here, we present the first integrated single-cell transcriptomic atlas of human cardiac organoids, combining 16 datasets generated across diverse differentiation strategies into a unified reference framework. The atlas resolves protocol-dependent effects on lineage composition, developmental fidelity, and cellular maturation, enabling direct comparison of cardiac organoid systems across the field. Mapping to fetal and adult heart references demonstrates that organoid-derived populations faithfully recapitulate early first-trimester developmental states, with maturation representing the major remaining bottleneck. The analysis identifies reproducible generation of epicardial and coronary vascular lineages, reveals distinct trade-offs between cellular diversity and maturation across protocols, and benchmarks organoids against complementary 3D cardiac models and newly generated datasets. Finally, drug-target analyses expose signaling pathways that remain insufficiently represented in vitro. This community resource provides a standardized framework for evaluating, comparing, and improving human cardiac differentiation models.

## Introduction

The human heart is a highly specialized multicellular organ that sustains the circulation of oxygen, nutrients and metabolic waste products throughout the body. Its function depends on the coordinated activity of different cell types that together govern electrical conduction, contractile force generation, vascular homeostasis and tissue remodeling [1][2]. The importance of this cellular coordination is underscored by the enormous global burden of heart disease. Cardiovascular disease remains a leading cause of morbidity and mortality worldwide [3][4], while congenital heart defects are the most common birth defects in humans [5]. Despite their diverse etiologies, both originate from perturbations of the cellular interactions, developmental programs and tissue architecture that underlie normal heart formation and function. Understanding these processes is therefore of fundamental biological and clinical importance.

Direct investigation of human cardiac development and disease remains challenging. Myocardial biopsies are not readily available and carry procedural risks [6], while post-mortem or transplant tissues are influenced by ischemia time, treatment history and terminal disease states and do not permit longitudinal analyses [7]. Furthermore, the earliest stages of human cardiogenesis, during which many congenital heart defects arise, are largely inaccessible. Although animal models have provided invaluable insights into cardiac biology, differences in developmental timing, physiology and therapeutic response can limit their ability to faithfully recapitulate human development and disease [8]-[11].

These limitations have driven the development of human pluripotent stem cell (hPSC)-derived cardiac models as experimentally tractable systems for studying human heart development, disease mechanisms and therapeutic interventions [12]-[25]. Such models span simple two-dimensional (2D) differentiation systems and increasingly complex three-dimensional (3D) platforms that recreate key features of cardiac tissue organization and cell-cell interactions. Heart organoids represent one prominent class of 3D model generating multiple cardiac lineages through directed differentiation and self-organization. Other approaches include cardiac micro-tissues assembled from separately differentiated cell types [25][26], embryoid bodies that undergo largely spontaneous differentiation, and engineered heart tissues, in which pre-differentiated cardiac cells are cultured with biomaterial scaffolds and exposed to mechanical and electrical conditioning to promote maturation [27]-[30].

For heart organoids, their utility ultimately depends on the extent to which they recapitulate the cellular, molecular, and developmental features of the human heart. During embryogenesis, cardiac progenitors arise from mesoderm and diversify into distinct progenitor populations that generate specialized cardiac lineages and structures [31]. The first and second heart fields give rise to complementary regions of the developing heart, with the former contributing primarily to the left ventricle and parts of the atria, and the latter generating the right ventricle, outflow tract and additional atrial and inflow tract components. Subsequently, the epicardium emerges from the proepicardium and produces much of the cardiac stroma and the mural cells of the coronary vasculature, while endothelial populations derived from the endocardium and sinus venosus assemble the developing coronary vascular network. As morphogenesis proceeds, cardiomyocytes, endothelial and endocardial cells, epicardial derivatives, and stromal populations engage in extensive reciprocal signaling that coordinates chamber patterning, myocardial growth, vascularization and functional maturation. Consequently, assessing whether heart organoids accurately reproduce the cellular diversity, developmental trajectories and intercellular communication networks of the developing human heart remains a central challenge for the field [32].

To model human cardiogenesis in vitro, heart organoid protocols attempt to recreate key developmental processes using a variety of differentiation strategies [12]-[25]. Most are based on the temporally controlled modulation of developmental signaling pathways, including canonical Wnt signaling, often combined with additional morphogens such as BMP, Activin A, FGF and retinoic acid [13]-[17][20]-[23][25][33][34]. Other approaches generate defined progenitor populations prior to 3D tissue formation or co-differentiate cardiac and non-cardiac lineages to capture specific developmental interactions [12][18][19][24]. However, protocols differ substantially in culture conditions, morphogen exposure, extracellular matrix requirements, and differentiation duration [12]-[25]. Whether and how these differences shape cellular composition, developmental trajectories, tissue architecture, and maturation state remains largely unresolved, limiting the comparison, standardization, and broader application of heart organoid models.

Single-cell transcriptomic techniques now offer an unprecedented opportunity to systematically address these questions. Large-scale initiatives such as the Human Cell Atlas have generated comprehensive reference maps of primary human tissues [35], including atlases of the developing and adult heart [2][36]. Similarly, integrated single-cell references have been established for organoids of the central nervous system and endoderm-derived organs, enabling robust comparisons across laboratories, differentiation protocols, and developmental stages [37]-[39]. No equivalent reference framework currently exists for heart organoids. As a result, datasets generated across studies remain difficult to compare, developmental trajectories cannot be readily aligned, and newly developed models lack a standardized framework for benchmarking against existing systems and the in vivo human heart.

Here, we present the Human Heart Organoid Cell Atlas (HHOCA), a comprehensive integrated reference of existing and unpublished single-cell transcriptomic datasets from hPSC-derived heart organoids, complemented by cardiac monolayer differentiation datasets as reference models. Using a harmonized annotation framework, we resolve cardiac cell states, lineages, and progenitor populations across multiple levels of biological organization. We define cardiomyocyte subtype programs, characterize intercellular communication networks and maturation trajectories, and benchmark organoid and monolayer-derived cell types against fetal and adult human heart references. We further perform a systematic cross-protocol comparison of cellular composition, lineage representation, and maturation states. Together, HHOCA establishes a comprehensive reference and benchmarking framework for heart organoids and related cardiac in vitro systems, enabling the rigorous assessment of their biological fidelity, experimental utility, and translational potential.

## Results

### HHOCA establishes a unified reference atlas of cardiac organoid development

To construct the HHOCA, we compiled single-cell transcriptomic data from 16 datasets including 14 publicly available and 2 unpublished, alongside 2D cardiac monolayer differentiation datasets as reference [12]-[25] (Fig. 1a). In total, the atlas comprises 422,059 cells, out of which 316,711 cells (∼75%) originate from organoids and 105,348 cells (∼25%) from monolayers, and encompasses 28 differentiation time points (Fig. 1b), providing broad coverage of cardiac lineage specification and maturation. Datasets span multiple stem cell lines and protocols, and were generated with different single-cell sequencing technologies, with most organoid datasets profiled using Chromium NextGem 3’ v3.1 (Fig. 1c, Table S1). To facilitate systematic comparison, protocols were grouped according to their developmental design logic into four classes (Table S1): 1) Wnt-modulation protocols, based on the temporal activation and inhibition of Wnt/beta-catenin signaling (GSK/Wnt inhibition; GiWi); 2) Developmental-morphogen protocols, using combinations of Wnt, BMP, Activin/Nodal, FGF, and sometimes retinoic-acid-pathway modulation (Multi-Factor); 3) Progenitor-expansion protocols, where cardiac mesoderm or cardiovascular progenitors are expanded before 3D organoid formation (Progenitor-Directed); and 4) Protocols directing co-development of cardiac and hematopoietic lineages (Co-Hematopoiesis).

**Figure 1:**
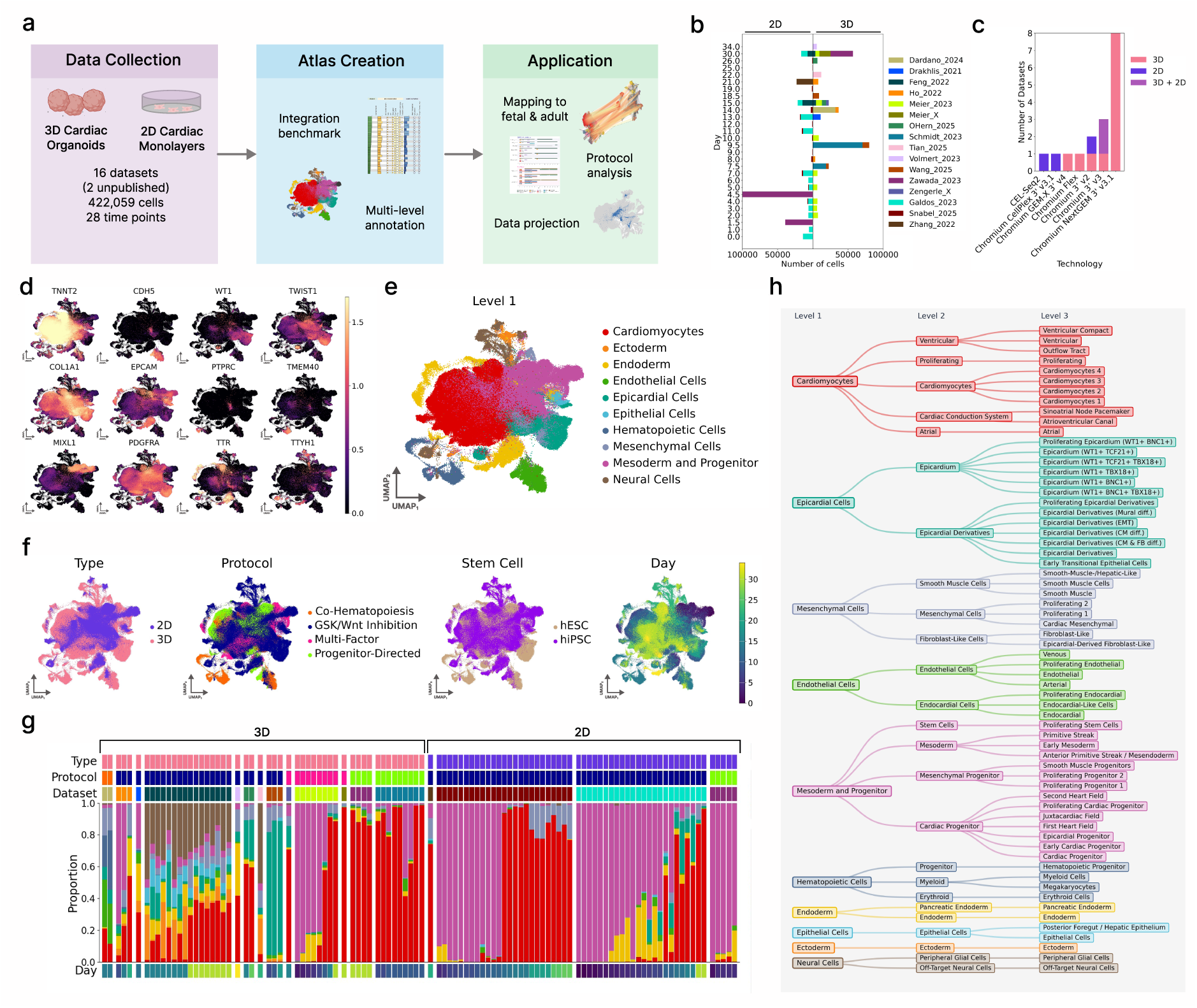
Integrated HHOCA reveals a multi-level cell type landscape of heart development in vitro. **a**, Overview of atlas construction. **b**, Number of cells per dataset across sampled time points (days), split by culture type: 2D (left) and 3D (right). **c,** Number of datasets per single-cell sequencing technology, colored by culture type (3D, 2D, or both). **d,** UMAP embeddings of the integrated atlas colored by expression of canonical marker genes. **e,** UMAP embedding of the integrated atlas colored by level 1 cell type annotation. **f,** UMAP embeddings of the integrated atlas colored by culture type (2D vs. 3D), protocol category, stem cell type (hESC vs. hiPSC), and sampling day. **g,** Proportion of level 1 cell types across all samples in the atlas, annotated by culture type, protocol, dataset (top tracks), and sampling day (bottom track). **h,** Hierarchical cell type annotation structure of level 1 (broad lineage), level 2 (cell type), and level 3 (fine-grained subtype/state) annotations across the atlas.

To create an integrated embedding, we benchmarked four methods using the single-cell atlassing toolbox [40], across three highly variable gene selections, two batch variables, and multiple hyperparameters (see Methods). Integration performance was assessed based on scIB metrics [41]. Because organoid and monolayer cultures differ substantially in cellular composition and developmental dynamics, we additionally generated separate integrated embeddings for each system, resulting in three complementary atlas representations (Fig. 1d-g, Fig. S1a-d). The final integrations enabled hierarchical cell type annotation at three levels of resolution (Fig. 1h, Fig. S1e,f). Cell type annotation was performed using a consensus gene marker list (Table S2) compiled from the 14 published heart organoid studies included in the atlas. At the broadest level, we identified 10 major cell types (Fig. 1d,e). Cardiomyocytes represented the largest population in organoids, followed by mesoderm and progenitor cells, epicardial cells, mesenchymal cells, and endothelial cells (Fig. S1a,g,h). In addition to cardiac lineages, organoids contained varying proportions of non-cardiac “off-target lineages”, including endodermal, neural, and ectodermal cells. Endodermal cells are an expected component of some protocols, as foregut endoderm co-develops with the heart in vivo and is deliberately co-induced in heart-forming organoids [13], whereas neural and ectodermal cells most likely reflect incomplete restriction of early differentiation towards mesoderm. Monolayer datasets were enriched for earlier developmental stages and were therefore dominated by mesodermal and progenitor populations, followed by cardiomyocytes (Fig. S1b,g,i). Smaller populations of epicardial and endothelial cells were also detected, together with off-target endodermal and neural cell types.

### Fine-grained annotation resolves cardiomyocyte, endothelial, epicardial, and heart field diversity

We next resolved the main cardiac lineages at higher resolution. Fine-grained annotation revealed a broad spectrum of cardiomyocyte, endothelial, and epicardial states, as well as distinct cardiac progenitor populations, capturing multiple developmental branches of human cardiogenesis (Fig. S2).

Cardiomyocytes yielded 11 transcriptionally distinct subclusters that could be grouped into 5 major categories: ventricular, atrial, cardiac conduction system, proliferating, and cardiomyocytes (Fig. 1h, Fig. S2a,b). Ventricular cardiomyocytes were characterized by expression of *MYH7* and *MYL2*, with ventricular compact cells additionally expressing *PRDM16* and *HEY2*. Cardiomyocytes of the outflow tract were identified by expression of *WNT5A*, *RSPO3*, and *FGF10*. Atrial cardiomyocytes expressed *NR2F2* and *KCNA5*, while specialized conduction system populations included sinoatrial node pacemaker cells (*SHOX2* and *HCN4*) and atrioventricular canal cells (*TBX2*, *TBX3*, *BMP2*, *RSPO3*, and *NR2F2*). Cardiomyocytes that did not show clear compartment-specific expression were annotated as “Cardiomyocytes”.

Endothelial cells comprised two broad categories: vascular endothelial cells and endocardial cells (Fig. 1h). Vascular endothelial cells further diversified into venous, arterial, proliferating, and unspecified endothelial (annotated as “Endothelial”) (Fig. S2c,d). Venous endothelial cells were characterized by expression of *EPHB4* and *NR2F2*, while arterial endothelial cells expressed the canonical arterial markers *CXCR4*, *SOX17*, *HEY2*, and *GJA4*. Endocardial cells consisted of three subpopulations: endocardial, endocardial-like, and proliferating endocardial (Fig. S2c,d). The endocardial subpopulation expressed established endocardial markers, including *NPR1*, *NFATC1*, and *NPR3*, together with the venous-associated genes *EPHB4* and *NR2F2*. In contrast, endocardial-like cells exhibited a hybrid transcriptional identity, co-expressing endothelial markers alongside cardiomyocyte genes such as *TNNT2* and *TTN*.

Cells of the epicardium displayed particularly pronounced heterogeneity and were categorized into two main groups: epicardial cells and epicardial derivatives (Fig. 1h). Distinct epicardial subtypes were defined by combinatorial expression of *WT1*, *BNC1*, *TBX18*, and *TCF21*, with individual populations distinguished by specific marker combinations (Fig. S2e,f). Epicardial derivatives were identified by either transcriptional signatures of epithelial-to-mesenchymal transition (EMT), marked by *TWIST1*, *SNAI1*, and *SNAI2,* or by lineage-specific differentiation programs (Fig. S2g). Among these, epicardial derivatives differentiating to cardiomyocyte/fibroblast (CM/FB diff.) expressed extracellular matrix and mesenchymal genes including *LUM*, *SEMA3D*, *TNC*, and *FN1*. A second population exhibited a cardiomyocyte-associated program (CM diff.), characterized by expression of *SFRP5* together with canonical cardiomyocyte markers such as *RYR2*, *TNNT2,* and *TTN*. Finally, mural-like epicardial derivatives (Mural diff.) expressed genes associated with mural commitment, including *PDGFRB* and *RGS5*. Together, these populations define a continuum of epicardial differentiation, from epicardial progenitor states through EMT-associated intermediates to lineage-committed derivatives, indicating that heart organoids recapitulate key aspects of epicardial maturation and fate diversification observed during heart development.

We also identified first heart field (FHF), second heart field (SHF), and juxtacardiac field (JCF) progenitors (Fig. S2h). FHF and SHF populations expressed shared early cardiac transcription factors such as *NKX2-5* and *MEF2C*, but presented distinct lineage signatures, with FHF cells expressing *HCN4* and *TBX5*, and SHF cells expressing *ISL1* and *FOXC1*. JCF progenitors were identified by the expression of *HAND1, MAB21L2, HOXB6, HOXB5,* and *BNC2*.

Collectively, these analyses show that heart organoids faithfully recapitulate key developmental trajectories of human cardiogenesis, generating a diverse repertoire of cardiac cell states encompassing heart field progenitors, specialized cardiomyocytes, and distinct endothelial and epicardial lineages.

### 3D tissue organization and signaling networks promote ventricular cardiomyocyte maturation

Beyond defining cellular composition, we asked how cardiac cell states evolve during differentiation and whether 2D and 3D cultures follow comparable developmental trajectories. To enable systematic comparisons, samples were grouped into three differentiation stages: early (day 0-10), middle (day 11-20), and late (day 21-35). This categorisation enabled direct comparison of the two culture formats at matched developmental stages while minimizing variation arising from differences in sampling time.

Among the annotated cell types, ventricular cardiomyocytes were the most abundant population in both monolayer and organoid cultures (Fig. S2i,j). Given their central role in cardiac tissue development and function, we further characterized this population. At the early stage, ventricular cardiomyocytes were less abundant in 2D monolayer cultures, precluding robust cross-format comparisons (Fig. 2a). Therefore, subsequent analyses focused on the middle and late stages, when ventricular cardiomyocytes were sufficiently represented in both systems.

**Figure 2:**
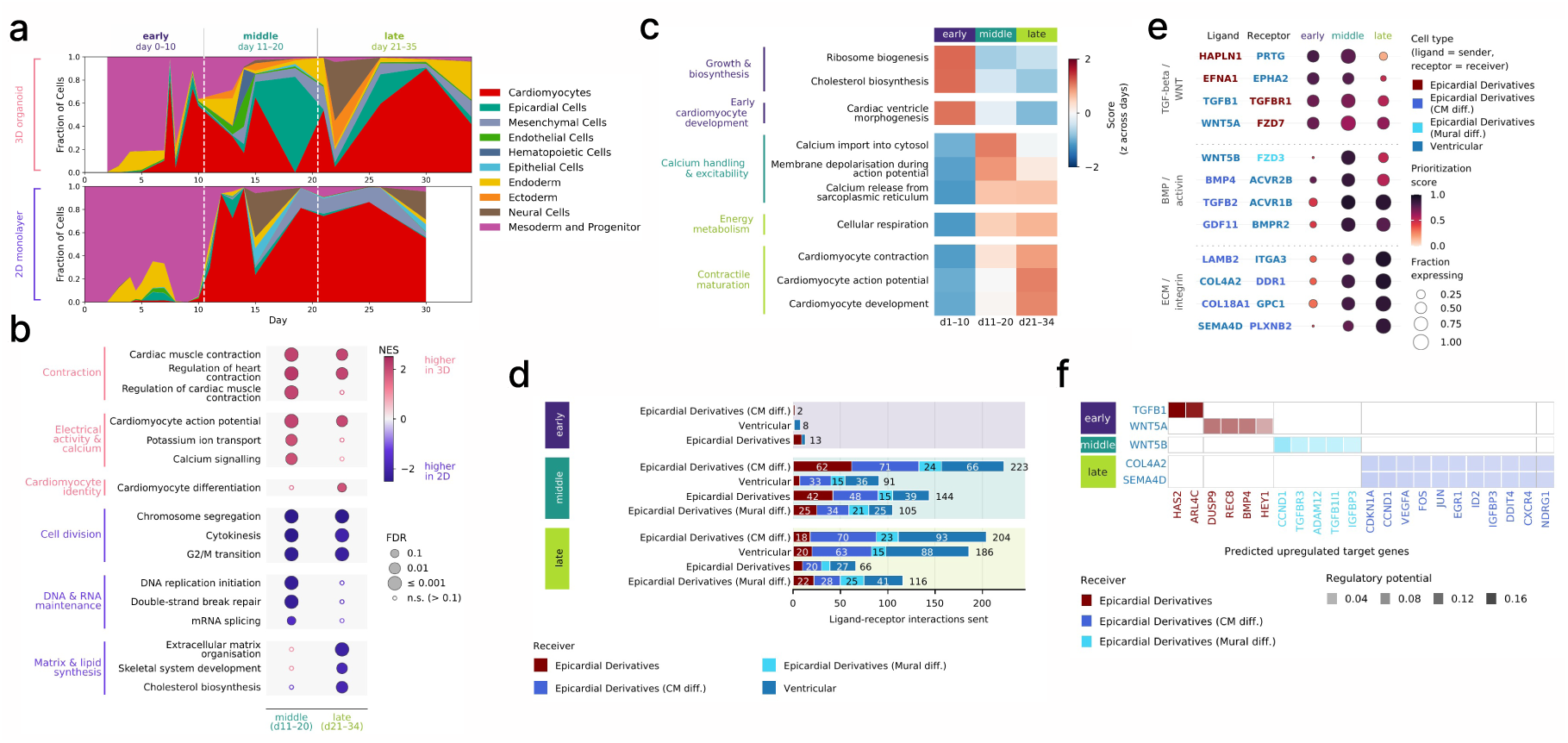
3D culture drives ventricular cardiomyocyte maturation and stage-specific communication with epicardial derivativesa, Cell-type composition per day of differentiation in 3D organoids (top; days 2-34) and 2D monolayers (bottom; days 0-30), shown as stacked fractions. Dashed lines separate the early (days 0–10), middle (days 11–20) and late (days 21–35) stages. **b,** GSEA of GO Biological Process terms comparing 2D and 3D ventricular cardiomyocytes at the middle (days 11-20) and late (days 21-35) stages. Terms enriched in the middle stage, late stage, or both stages are shown. color indicates normalized enrichment score (NES), with negative values indicating enrichment in 2D and positive values indicating enrichment in 3D. Dot size indicates FDR; terms with FDR ≤ 0.05 are shown. **c,** Activity of selected GO Biological Process programs in 3D ventricular cardiomyocytes across differentiation windows (days 0-10, 11-20 and 21-35). NES values were z-scored within each window for each program. The top bar indicates the differentiation stage. **d,** MultiNicheNet-inferred cell-cell communication between ventricular cardiomyocytes and co-occurring cell types in 3D organoids at each stage. Chord width indicates the number of inferred ligand-receptor interactions between cell types; the total number of interactions is shown above each plot. **e,** Top-prioritized ligand-receptor pairs involving ventricular cardiomyocytes, grouped by the stage at which each pair received its highest prioritization score and shown across all stages. Ligand and receptor names are colored according to the cell type expressing each gene. Dot color indicates the MultiNicheNet prioritization score and dot size indicates the fraction of cells expressing the gene. **f,** Predicted target genes of ligands secreted by ventricular cardiomyocytes in receiver populations, grouped by stage. Target genes are colored according to receiver cell type, and tile shading indicates ligand-target regulatory potential.

Differential expression analysis, followed by Gene Ontology Biological Process enrichment of ranked genes, identified consistent differences between culture formats. At both intermediate and late stages, ventricular cardiomyocytes from monolayer culture were enriched for proliferative programs, including G2/M phase transition, mitotic cytokinesis and sister chromatid segregation, and DNA replication initiation and homologous recombination repair at the middle stage (Fig. 2b). In contrast, organoid-derived ventricular cardiomyocytes preferentially expressed pathways associated with cardiac function and maturation, including muscle contraction, cardiac conduction, action potential, and ventricular cardiac muscle tissue development (Fig. 2b). During normal heart development in vivo, ventricular cardiomyocytes progressively reduce proliferation while acquiring specialized contractile, electrophysiological and metabolic features [42]. The persistence of proliferative programs in 2D cultures, together with the enrichment of cardiac functional programs in organoids at matched developmental stages aligns with the notion that the 3D multicellular environment, with its cell-cell interactions, mechanical cues and tissue-like architecture, actively supports cardiomyocyte maturation and functional specialization.

To further characterize ventricular cardiomyocyte maturation within organoids, we examined transcriptional programs across the full differentiation time course (Fig. 2c). Early cardiomyocytes (days 0-10) were dominated by biosynthetic and morphogenetic programs, including ribosome biogenesis, cholesterol biosynthesis and cardiac ventricle morphogenesis. During the intermediate stage (days 11-20), these programs declined, coinciding with increased expression of pathways involved in calcium handling and electrical excitability. By the late stage (days 21-35), cardiomyocyte development, action potential, contraction and cellular respiration programs reached the highest levels. Ventricular cardiomyocytes thus progressed from a growth-oriented biosynthetic state toward electrophysiological, contractile and oxidative metabolic maturation, closely paralleling key features of in vivo cardiogenesis [42].

Given that cardiomyocyte maturation occurs within a multicellular tissue environment involving dynamic interactions with endothelial and epicardial lineages, we next sought to define how intercellular communication networks evolve during organoid development. To this end, we applied MultiNicheNet [43] across the three differentiation stages. Because endothelial cells were represented in fewer samples than required for robust MultiNicheNet analysis (see Methods), we focused on signaling interactions among ventricular cardiomyocytes and epicardial derivatives, including the CM- and mural-differentiating subsets. The complexity of predicted communication networks increased substantially during differentiation, from 23 interactions at the early stage to 563 at the intermediate stage and 572 at the late stage (Fig. 2d), indicating progressive establishment of multicellular signaling networks during organoid maturation. As ventricular cardiomyocytes matured, epicardial-derived matrix and contact cues to ventricular cardiomyocytes, including *HAPLN1-PRTG* and *EFNA1-EPHA2*, progressively declined. This is consistent with a role for the epicardial niche in supporting myocardial expansion, as *HAPLN1*-expressing epicardial cells have previously been implicated in cardiomyocyte proliferation and ventricular-wall maturation [44]. In contrast, predicted cardiomyocyte-derived TGFB1 and *WNT5A* signaling to epicardial derivatives, linked to epicardial EMT and non-canonical Wnt signaling in cardiogenesis [45], persisted throughout (Fig. 2e). Stage-specific signaling programs emerged along epicardial differentiation trajectories. At the middle stage, the “CM-diff” epicardial branch was predicted to signal to ventricular cardiomyocytes through multiple TGF-β-family ligands acting on shared activin/BMP receptors (*TGFB2-ACVR1B, GDF11-BMPR2, BMP4-ACVR2B*). These pathway-level predictions are consistent with known roles for myocardial TGF-β2 in ventricular wall development [46] and for activin type II receptor ligands in cardiomyocyte proliferation [47]. Ventricular cardiomyocytes also exhibited reciprocal communication with mural-derived epicardial cells (Mural diff) via the non-canonical *WNT5-FZD36* axis, suggesting a potential role for myocardial signaling in mural-cell organization and vascular development. At the late stage, communication shifted towards extracellular-matrix-associated interactions. Epicardial derivatives predominantly expressed *LAMB2* and *COL18A1*, while cardiomyocytes *COL4A2*. Both populations also expressed receptors for matrix components generated by the other cell type, suggesting potential reciprocal matrix-mediated interactions. Given that laminin has been implicated in cardiomyocyte maturation [48] and the epicardium deposits a laminin sheath around the ventricle [49], this suggests a cooperative assembly of a matrix niche supporting ventricular cardiomyocyte maturation. Analysis of predicted downstream target genes further supported these stage-specific programs (Fig. 2f). Early ventricular cardiomyocyte-derived *TGFB1* and *WNT5A* were associated with expression of *HAS2, BMP4* and *HEY1* in epicardial derivatives, consistent with epicardial activation and early extracellular-matrix production. In contrast, at the middle stage, *WNT5B* was predicted to upregulate proliferative and mural-like genes (*CCND1, TGFBR3, ADAM12, TGFB1I1, IGFBP3*) specifically in mural-differentiating epicardial derivatives. At the late-stage, *COL4A2* and *SEMA4D* were associated with a shared set of immediate-early and stress-response genes (*FOS, JUN, EGR1, CDKN1A, DDIT4*) as well as *VEGFA* and *CXCR4* in CM-differentiating epicardial derivatives. Overall, the predicted targets suggest that ventricular cardiomyocyte signaling shifts during development from activating early epicardial derivatives, through branch-specific instruction of the mural lineage, towards extracellular matrix-related signaling at later stages. The analysis also identifies candidate interactions that may further inform cardiomyocyte maturation (Fig. 2f).

### Heart organoids recapitulate fetal cardiac lineages but remain developmentally immature

To assess how well in vitro models can recapitulate in vivo transcriptional profiles, we compared the HHOCA with single-cell atlases of the developing human heart [36] and adult human heart [2]. We performed neighborhood matching between the atlases using scRabbit [50] and evaluated to which extent organoid and monolayer cell types correspond to their in vivo counterparts (see Methods). Broad cellular identities were successfully matched when organoids were compared with fetal heart data, particularly for cardiomyocytes and endothelial cells (Fig. 3a,b). Epicardial cells showed preferential matching to fetal mesenchymal populations, possibly reflecting the large proportion of epicardial cells undergoing EMT in organoids, while neural off-target cells matched fetal neural cells, further confirming the accuracy of the annotation. Similar correspondence was observed for monolayer-derived cell types (Fig. S3a,b). In contrast, mapping to the adult heart was substantially weaker, despite retention of broad lineage identities (Fig. S3c-f). The reduced correspondence to adult cardiac cell types was particularly apparent in monolayer cultures, where even cardiomyocytes exhibited only limited matching to adult counterparts. These findings suggest that although lineage specification occurs efficiently in both systems, acquisition of terminally differentiated cardiac states remains incomplete. The stronger correspondence observed in organoids further suggests that multicellular three-dimensional tissue organization contributes to developmental fidelity beyond lineage specification alone.

**Figure 3:**
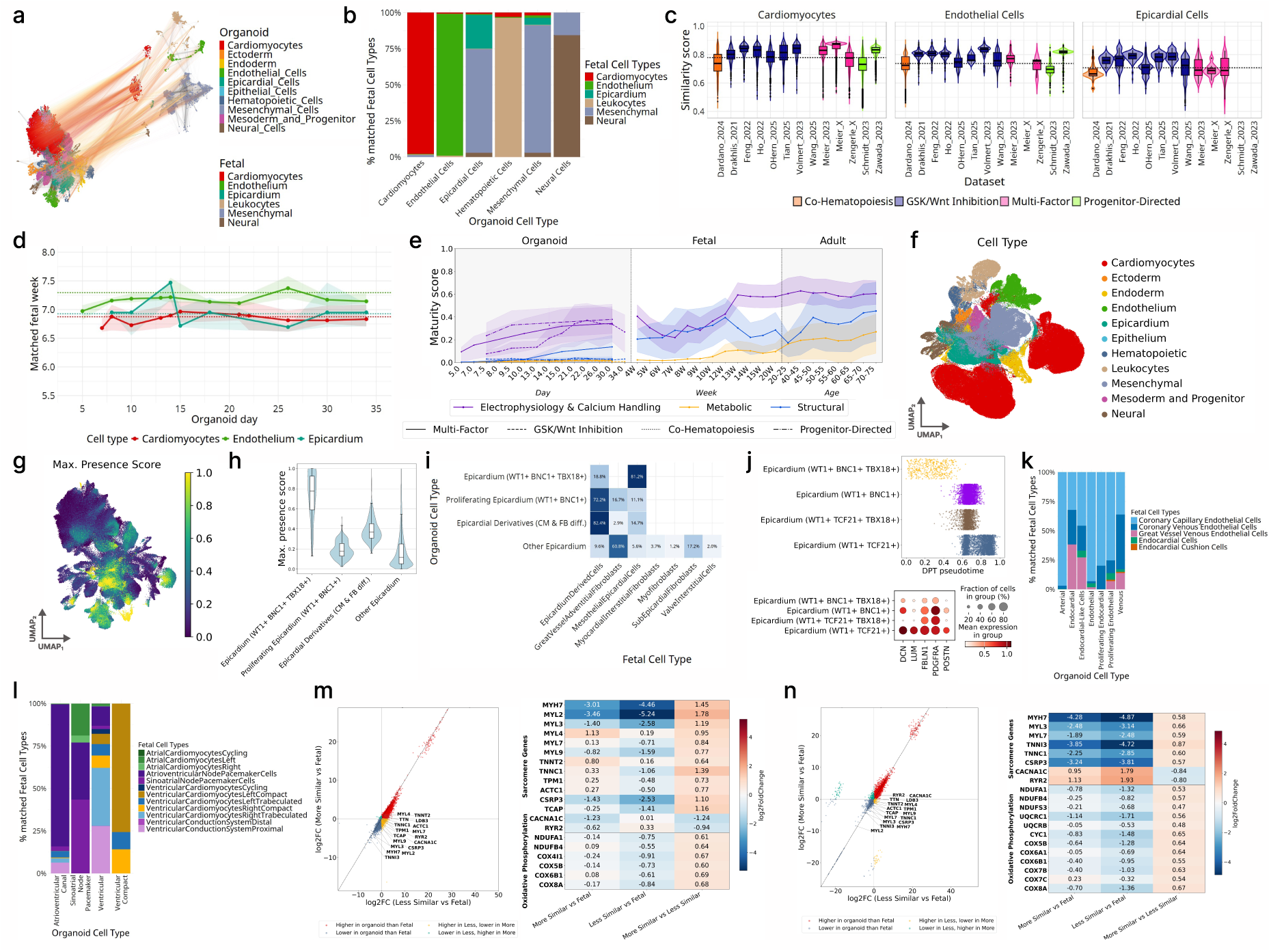
Fetal mapping reveals protocol-dependent maturation and subtype-specific fidelity across cardiomyocyte, endothelial, and epicardial lineages a, Unsupervised neighborhood matching between the 3D organoid atlas (left) and a fetal heart reference atlas (right) using scRabbit. Lines connect organoid and fetal neighborhoods identified as mutual nearest neighbors in a shared expression space, colored by level 1 cell type. **b,** Proportion of matched fetal cell types for each organoid cell type, showing the correspondence between level 1 organoid and broad fetal cell type annotations. **c,** Similarity scores between organoid and fetal cardiomyocytes, endothelium, and epicardium across datasets, colored by protocol type. The dotted lines show the mean. **d,** Matched fetal week for organoid cardiomyocytes, endothelium, and epicardium across organoid day. The dotted lines show the mean. **e,** Maturity scores (electrophysiology and calcium handling, metabolic, and structural; see Methods) across organoid, fetal, and adult developmental time. Organoid maturity scores are stratified by protocol type. **f,** UMAP embedding of the integrated organoid and fetal atlases, colored by cell type. **g,** UMAP embedding of the integrated organoid atlas colored by maximum presence score (see Methods). **h,** Violin plots of maximum presence scores for epicardial subtypes. **i,** Heatmap showing the percentage of matched cells between organoid epicardial subtypes and fetal epicardial/mesenchymal cell types. **j,** Diffusion pseudotime ordering of epicardial subtypes (top) and dot plot of fibroblast commitment marker expression across epicardial subtypes (bottom). **k,** Proportion of matched fetal endothelial cell types for each organoid endothelial cell type **l,** Proportion of matched fetal cardiomyocyte subtypes for organoid cardiomyocytes. **m,** Convergence of organoid ventricular cardiomyocyte gene expression toward fetal expression. Left, scatter plot comparing log2 fold change (organoid vs. fetal) between more similar and less similar organoid cells. Right, heatmap of log2 fold change for sarcomere and oxidative phosphorylation genes across comparisons. **n,** As in **m**, for organoid compact cardiomyocytes.

We next asked whether developmental fidelity varies across organoid-generating strategies. To this end, we examined neighborhood similarity scores for cardiomyocytes, endothelial cells, and epicardial cells that successfully matched their fetal counterparts (Fig. 3c). The similarity scores quantify the Spearman correlation of the gene expression between the matched neighborhoods [50]. On average, organoid cardiomyocytes and endothelial cells showed higher similarity to fetal reference populations than epicardial cells. Among protocols, GiWi and Multi-Factor approaches generally achieved the highest similarity scores. We further estimated the developmental stage represented by organoid-derived cell types. Across lineages, endothelial cells, cardiomyocytes, and epicardial cells most frequently mapped to fetal hearts between 6.5 and 7.5 weeks of development, with endothelial populations exhibiting the most advanced developmental profiles (Fig. 3d).

To quantify cardiomyocyte maturation more directly, we computed gene-set-based maturity scores reflecting electrophysiology and calcium handling, metabolic, and structural maturation programs across organoids, fetal, and adult cardiomyocytes (Fig. 3e; see Methods). The maturity scores correspond to gene scores computed using the pyUCell package [51] of maturity genes curated from literature (Table S3). Metabolic maturation remained largely absent in organoid datasets and increased substantially only at later fetal stages and in adult tissue. This is consistent with the well-described persistence of glycolytic metabolism in hPSC-derived cardiomyocytes and the postnatal transition towards fatty acid oxidation in vivo [42]. In contrast, electrophysiological and structural maturation varied considerably between protocols. Notably, Multi-Factor protocols consistently exhibited higher electrophysiological and calcium-handling, and structural maturity scores than GiWi-based approaches (Fig. 3e). These findings indicate that differences in protocol design influence the extent of cardiomyocyte maturation even when lineage specification is broadly preserved.

### Cell-state-level comparison reveals lineage-specific differences in developmental fidelity

While broad lineage corresponded well to fetal cardiac populations, developmental fidelity may vary considerably among individual cell states within each lineage. We therefore next assessed developmental fidelity at higher resolution by comparing organoid and monolayer subpopulations directly with their in vivo fetal counterparts. To this end, we integrated in vitro and in vivo datasets using sysVI [52], generating shared embeddings that enabled comparison of individual cell states across systems (Fig. 3f, Fig. S3g). Based on these embeddings, we computed presence scores [38] of organoid and monolayer cells, which quantify the likelihood of a given in vitro cell state appearing in the corresponding in vivo data (Fig. 3g).

Combining the neighborhood matching with the presence score revealed substantial differences in fidelity between individual cell states. Within the epicardial compartment, the *WT1+BNC1+TBX18+* epicardial subtype showed the strongest correspondence to fetal mesothelial epicardium (Fig. 3h,i). Among epicardial derivatives, only those with CM/FB differentiation closely matched their fetal counterparts, while the remaining epicardial subtypes and derivatives largely mapped to great vessel adventitial fibroblasts and showed comparatively low presence scores. Pseudotime and marker genes analysis further suggested that *WT1+BNC1+* and *WT1+TCF21+TBX18+* populations are transitional states progressing towards epicardial-derived cells, while *WT1+TCF21+* epicardial cells appear committed towards a fibroblast fate (Fig. 3j). Interestingly, epicardial cells derived from monolayers also showed partial correspondence to fetal epicardium (Fig. S3h,i).

Within the endothelial lineage, both organoid and monolayer cells predominantly mapped to the fetal coronary capillary endothelial population (Fig. 3k, Fig. S3j). Although organoid arterial endothelial cells expressed canonical arterial specification markers (*GJA4, SOX17, CXCR4, HEY2*), they yet failed to recapitulate the complete fetal arterial endothelial transcriptome, likely reflecting an early arterial identity that lacks the maturation signals present in the developing heart. In contrast, venous endothelial cells showed the strongest correspondence to their fetal counterpart, with approximately 60% of cells successfully matching. Endocardial and endocardial-like cells mapped predominantly to great vessel and coronary venous endothelial cells, as well as coronary capillary endothelial cells. Together, these findings suggest that endothelial diversification is readily achieved in organoids, whereas arterial and endocardial specialization may remain incomplete.

Focusing on cardiomyocytes, organoid ventricular cells robustly matched to their fetal counterparts (Fig. 3l). Other successful matches included organoid atrioventricular canal cells to atrioventricular node pacemaker cells. Sinoatrial node pacemaker cells mapped preferentially to their fetal counterpart, while also showing partial transcriptional overlap with atrioventricular-node pacemaker cells and atrial cardiomyocytes, aligned with their shared conduction-system and atrial developmental origins. Notably, both organoid and monolayer compact cardiomyocytes showed strong correspondence to fetal left ventricular compact cardiomyocytes, a result further supported by their high presence scores (Fig. 3l, Fig. S3h,k). When examining organoid and monolayer cardiomyocytes that successfully mapped to adult cardiomyocytes, they matched specifically to adult atrial cardiomyocytes (Fig. S3d,f), consistent with previous observations that immature ventricular cardiomyocytes share transcriptional features with mature atrial cardiomyocytes [53][54].

We next sought to determine whether reduced fidelity reflected incorrect lineage specification or incomplete maturation. To address this, we compared ventricular and compact cardiomyocytes from organoids with their fetal counterparts, stratifying organoid cells into less similar (similarity score < 0.5) and more similar (similarity score > 0.5) groups (Fig. 3m,n). Relative to their fetal cardiomyocytes, organoid cells consistently exhibited reduced expression of sarcomere genes, such as *MYH7* and *MYL3*, and oxidative phosphorylation genes. Importantly, these deficiencies were less pronounced in the more similar group, indicating that the primary distinction between less and more fetal-like cardiomyocytes lies in the degree of maturation achieved rather than in lineage identity. Thus, incomplete maturation, rather than incorrect specification, appears to be the main factor limiting the fidelity of organoid ventricular cardiomyocytes.

### Protocol design shapes the balance between cellular diversity and maturation

Having established that incomplete maturation is a major factor limiting organoid fidelity, we next asked how protocol design influences cellular composition and developmental state. For this analysis, we focused on organoid datasets that aimed to generate organized, comprehensive cardiac structures starting from hPSCs (GiWi, Multi-Factor, Co-Hematopoiesis) rather than cell populations from distinct cardiac progenitors obtained in 2D (Progenitor-Directed). Although protocols within each class also differed in morphogen concentrations, timing, and stem cell lines (Fig. 4a,b), this framework enabled systematic evaluation of the principal design strategies currently used to generate heart organoids.

**Figure 4:**
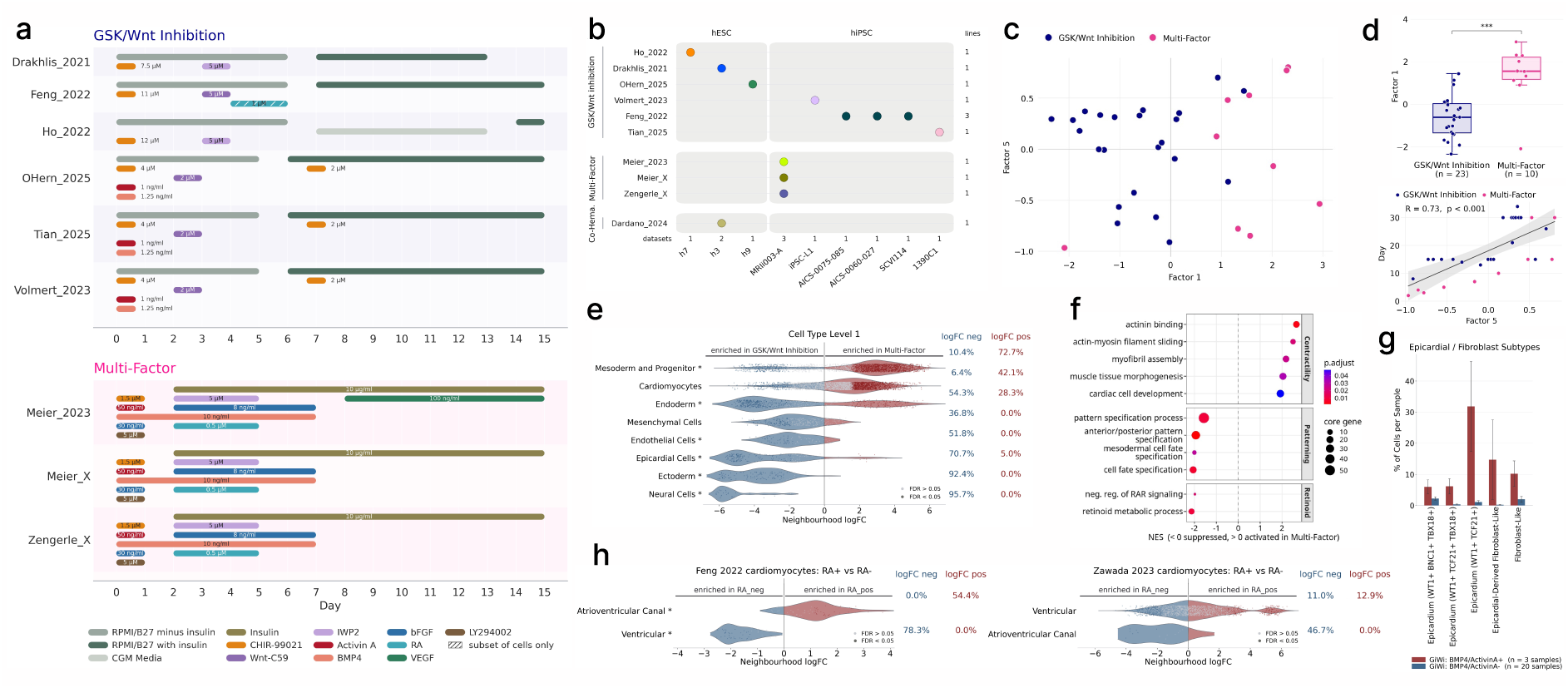
GSK/Wnt inhibition and Multi-Factor protocols yield distinct cell type compositions and pathway activity a, Overview of media composition and morphogen concentrations over culture time for organoid GiWi (top) and Multi-Factor (bottom) protocols, colored by supplement. **b,** Stem cell lines used by each dataset, split by hESC and hiPSC and protocol type, colored by dataset; the number of lines used per dataset is indicated on the right. The number of times a line is used by datasets at the bottom. **c,** MOFA embedding of samples based on Factor 1 and Factor 5. **d,** Factor 1 scores compared between GiWi and Multi-Factor protocols, showing significantly higher scores in Multi-Factor samples (top; *** p < 0.001), and Factor 5 scores plotted against differentiation day, showing a positive correlation (bottom; R = 0.73, p < 0.001). **e,** Differential abundance of level 1 cell types between GiWi and Multi-Factor protocols, tested using Milo. Points represent individual neighborhoods, colored by significance (FDR < 0.05). Percentages of significant neighborhoods enriched in each direction are shown on the right. A star indicates that at least 50% of neighborhoods are significant in the same direction. **f,** Gene set enrichment analysis of genes activated or suppressed in Multi-Factor relative to GiWi protocols, grouped by biological theme (contractility, patterning, retinoid). **g,** Percentage of epicardial and fibroblast subtypes per sample, compared between BMP4/Activin A+ and BMP4/Activin A-GiWi organoids. **h,** Differential abundance of cardiomyocyte subtypes between RA+ and RA-conditions for Feng et al. (left) and Zawada et al. (right), tested using Milo. Points represent individual neighborhoods, colored by significance (FDR < 0.05). Percentages of significant neighborhoods enriched in each direction are shown on the right. A star indicates that at least 50% of neighborhoods are significant in the same direction

To identify the main sources of variation across samples, we applied Multi-Omics Factor Analysis (MOFA) [55] to GiWi (n = 23) and Multi-Factor (n = 10) organoid samples, treating each cell type-specific transcriptome and cell-state composition as a separate view (see Methods). MOFA inferred 10 latent factors, of which two captured the dominant biological variation across the atlas (Fig. 4c,d). Factor 1 primarily separated samples according to protocol type, with Multi-Factor samples exhibiting significantly higher values than GiWi samples (Wilcoxon rank-sum test, p < 0.001; Fig. 4c,d). Because protocol type is partially confounded by study of origin, protocol-specific effects cannot be fully disentangled from study-specific technical variation; nevertheless factor 1 accounted for most of the variance within mesenchymal (∼65%) and epicardial (∼50%) populations (Fig. S4a). Its top-weighted features came almost exclusively from the mesenchymal view. They were dominated by genes involved in oxidative phosphorylation (e.g., NDUFB family, *COX4I1, COX7C, CYCS*), proteasomal degradation (*PSMA7, PSMB1*) and translation (*EIF2S2, EIF3I*), all of which were consistently higher in Multi-Factor samples (Fig. S4b,c). In contrast, Factor 5 captured variation shared across all cell types and correlated strongly with differentiation time (Pearson R = 0.73, p < 0.001; Fig. 4d), but showed no association with protocol type (Fig. 4d). Together, these analyses indicate that protocol design and developmental stage represent distinct sources of variation across heart organoid datasets. Notably, we observed no clear differences associated with the hPSC line (Fig. S4d) suggesting that protocol-associated effects outweigh differences attributable to genetic background in the datasets considered.

#### Multi-Factor and GiWi protocols favor distinct cellular and maturation outcomes

We next examined whether these protocol-associated sources of variation were accompanied by changes in cellular composition. Differential abundance analysis using Milo [56] revealed striking differences between GiWi and Multi-Factor organoids (Fig. 4e). Multi-Factor organoids were dominated by cardiomyocytes, whereas GiWi organoids displayed substantially greater cellular diversity, including increased representation of endothelial, mesenchymal, and epicardial populations (Fig. 4e). Off-target lineages, particularly ectoderm and neural populations, were almost exclusively present in GiWi organoids. Thus, while Multi-Factor protocols favor cardiomyocyte generation, GiWi protocols produce a broader spectrum of cardiac and non-cardiac cell types.

To determine whether protocol design also influences cardiomyocyte state, we compared ventricular cardiomyocyte transcriptional programs between protocol classes (Fig. 4f). Gene Set Enrichment Analysis revealed enrichment of contractility-associated terms in Multi-Factor cardiomyocytes, while GiWi-derived cardiomyocytes preferentially expressed developmental patterning and retinoid-related programs (Fig. 4f). These findings are consistent with the increased maturity of Multi-Factor cardiomyocytes observed in the fetal comparison analyses (Fig. 3).

#### Morphogen supplementation directs lineage diversification within GiWi organoids

To further dissect the sources of variation within GiWi protocols, we evaluated the influence of BMP4 and Activin A supplementation using the same Milo differential abundance framework. We excluded the Tian_2025 dataset [21], which was dominated by ectodermal and neural cells and therefore not suitable for this comparison. Differential abundance analysis revealed that BMP4/ActivinA supplementation profoundly affected epicardial and fibroblast-related populations (Fig. 4g, Fig. S4e). BMP4/ActivinA-positive protocols showed significantly higher abundance of fibroblast-like and epicardial-derived fibroblast-like cells, as well as the *WT1+TCF21+* epicardial state (Fig. 4g, Fig. S4e). Notably, these protocols also enriched for the *WT1+BNC1+TBX18+* epicardial population, which most closely resembled fetal epicardium in our comparison analyses, and for the transitional *WT1+TCF21+TBX18+* epicardial state. Within the cardiomyocyte compartment, BMP4/ActivinA supplementation increased the abundance of sinoatrial node pacemaker cells, and ventricular compact cardiomyocytes (Fig. S4f). Transcriptionally, ventricular cardiomyocytes from BMP4/ActivinA+ organoids were enriched for ion channel-related pathways, while BMP4/ActivinA-negative conditions showed stronger enrichment of processes related to oxidative phosphorylation (Fig. S4g).

#### Retinoic acid effects depend on differentiation context

Finally, we explored the influence of retinoic acid (RA) using two studies that each compared RA-positive and RA-negative conditions, while noting that the concentration and timing of RA exposure differed substantially between studies [16][24]. In the Feng dataset [16], RA withdrawal favored ventricular cardiomyocyte formation, whereas RA supplementation increased the abundance of atrioventricular canal cells (Fig. 4h), consistent with known roles of RA signaling in cardiac patterning [57][58]. In contrast, RA treatment had comparatively modest effects in the Zawada dataset [24], where ventricular cardiomyocyte abundance remained largely unchanged between conditions. As these studies differed in protocol design, RA dosage, and timing of exposure, the observed effects cannot be attributed solely to RA signaling, but instead suggest that the impact of RA is shaped by its interaction with the broader differentiation context.

Collectively, these analyses demonstrate that protocol design is a major determinant of both cellular composition and developmental state in heart organoids. Multi-Factor protocols preferentially generate cardiomyocyte-rich organoids with more mature transcriptional programs, whereas GiWi protocols produce more heterogeneous tissues containing a broader repertoire of cardiac and non-cardiac lineages. Within GiWi protocols, BMP4 and Activin A supplementation promotes the generation of epicardial and fibroblast-associated populations. We also observed differing effects of RA supplementation, depending on the specific dataset.

### Atlas mapping enables benchmarking of diverse cardiac model systems

A major goal of the HHOCA is to provide a common reference framework for comparing diverse cardiac in vitro systems and for benchmarking newly generated datasets. To test this capability, we projected cardiac micro-tissues [25], cardiac-directed embryoid bodies [20], and an independent heart organoid dataset [59] onto the atlas using scArches [60].

Cardiac micro-tissues are generated by first differentiating individual cell types, such as cardiomyocytes, mesenchymal cells, and endothelial cells, and subsequently assembling them into a 3D tissue construct (Fig. 5a). To compare micro-tissues with self-organizing organoids, we mapped a day 34 micro-tissue dataset [25] onto the 3D atlas and trained a custom CellTypist [61] model on the level 1 annotation of the atlas to predict micro-tissue cell types (Fig. 5b,c). Overall, cardiomyocytes and mesenchymal cells from micro-tissues showed low mapping quality scores [62] (Fig. 5d), indicating transcriptional states that differed substantially from those observed in organoids. Gene set enrichment analysis revealed increased metabolic maturity in both micro-tissue cardiomyocytes and mesenchymal cells, whereas organoid cells were enriched for fetal developmental programs (Fig. 5e, Fig. S5a). In contrast, electrophysiology and calcium handling, and structural maturity scores were higher in organoid cardiomyocytes than in stage-matched micro-tissues (Fig. S5b). These observations suggest that engineered assembly promotes metabolic maturation, whereas the self-organized co-development of multiple cardiac lineages favors the acquisition of developmental, structural and electrophysiological programs.

**Figure 5:**
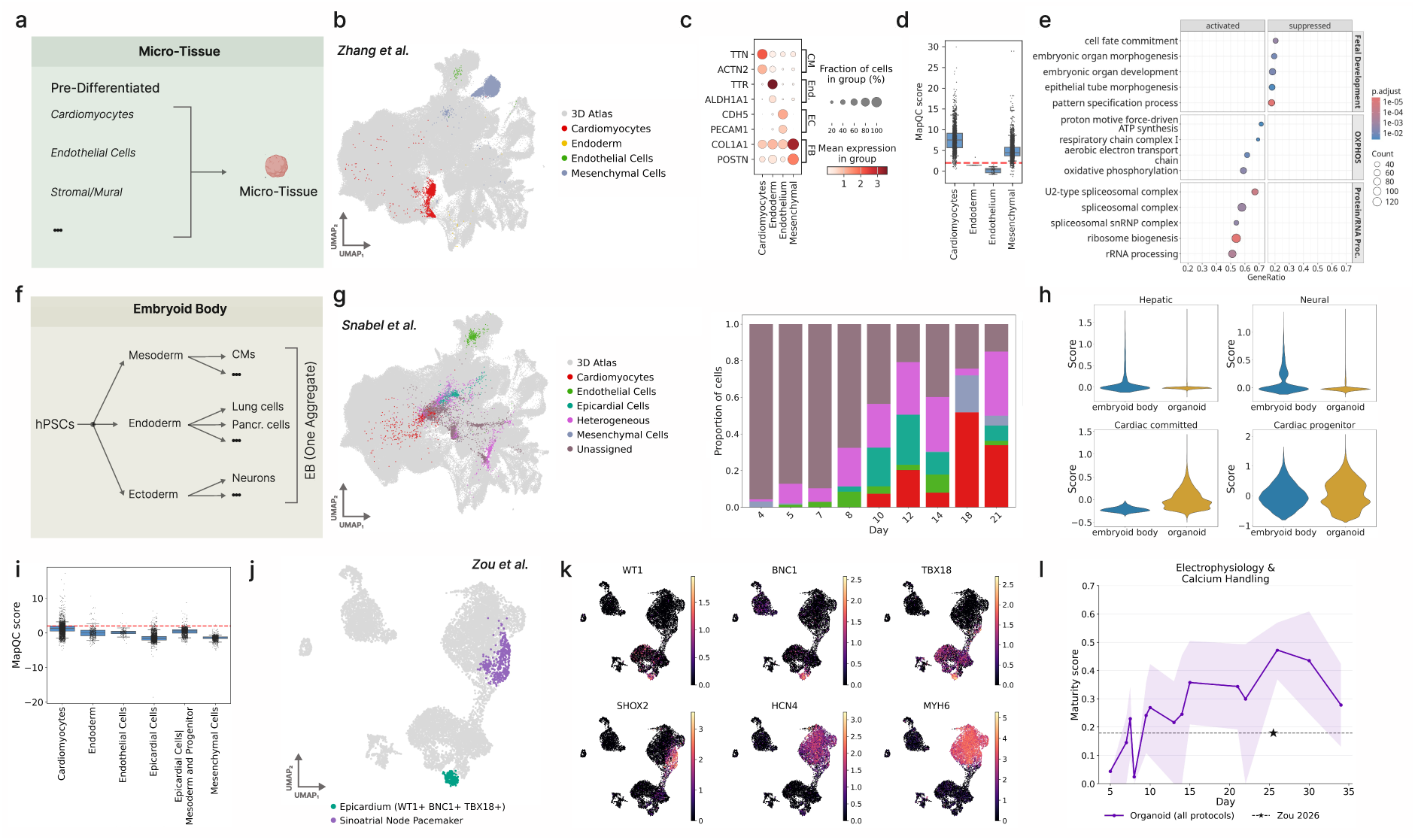
Reference mapping reveals cell identity and maturity differences across micro-tissues, embryoid bodies, and a query organoid dataset a, Schematic of micro-tissue generation (“…” indicates further cell types). **b,** UMAP embedding of the 3D organoid atlas (grey; same embedding as in Fig. S1) with micro-tissue cells [25] mapped on, colored by predicted cell type. **c,** Dot plot of marker gene expression for cardiomyocytes (CM), endoderm (End.), endothelial cells (EC), and fibroblasts (FB) of the predicted micro-tissue cell types. **d,** MapQC scores for micro-tissue cardiomyocytes, endoderm, endothelium, and mesenchymal cells, indicating mapping quality relative to the organoid atlas; dashed line indicates the quality threshold (MapQC score = 2). **e,** Gene set enrichment analysis of pathways activated or suppressed in micro-tissue cardiomyocytes relative to organoids, grouped by biological theme (fetal development, OXPHOS, protein/RNA processing). **f,** Schematic of embryoid body generation (“…” indicates further cell types). **g,** UMAP embedding of the 3D organoid atlas (grey; same embedding as in Fig. S1) with embryoid body cells [20] mapped on, colored by predicted cell type (left), and proportion of predicted cell types in embryoid bodies across sampled days (right). **h,** Violin plots comparing hepatic, neural, cardiac-committed, and cardiac progenitor gene scores between embryoid body and organoid cells **i,** MapQC scores for cell types of the query organoid dataset [59] mapped onto the atlas; dashed line indicates the quality threshold (MapQC score = 2). **j,** UMAP embedding of the query organoid cells, highlighting WT1+BNC1+TBX18+ epicardium and sinoatrial node pacemaker cells. **k,** UMAP embeddings of the query organoid cells colored by expression of epicardial (WT1, BNC1, TBX18), sinoatrial node pacemaker (SHOX2, HCN4) and cardiomyocyte (MYH6) marker genes. **l,** Electrophysiology and calcium handling maturity scores of cardiomyocytes(see Methods) across organoid day, comparing all organoid protocols in the atlas (blue line) to the query organoid dataset (black star).

We next asked how cardiac-directed embryoid bodies relate to heart organoids. Unlike organoids, embryoid bodies arise through spontaneous differentiation and contain derivatives of all three germ layers, although their differentiation can be biased toward cardiac fates through directed signaling cues (Fig. 5f) [20][27][28]. We mapped a cardiac-directed embryoid body dataset spanning 9 time points [20] onto the 3D atlas and predicted cell identities using the same CellTypist classifier (Fig. 5g, Fig. S5c). Early embryoid body populations showed poor correspondence to organoid progenitors and were mostly classified as unassigned (Fig. 5g), reflecting a differentiation trajectory distinct from that present in cardiac organoids. Even at later stages, many cells remained heterogeneous or unassigned, likely reflecting populations absent from cardiac organoids. Consistent with this interpretation, embryoid body progenitors retained transcriptional signatures associated with hepatic, neural, and endodermal lineages, whereas organoid progenitors were strongly enriched for cardiac commitment programs (Fig. 5h). Despite these divergent developmental routes, embryoid body-derived cardiomyocytes ultimately achieved electrophysiological and structural maturity comparable to organoid cardiomyocytes, while exhibiting slightly higher metabolic maturity (Fig. S5d).

A key application of a reference atlas is the rapid, standardized annotation and benchmarking of newly generated datasets, without requiring manual cell type annotation. To demonstrate this, we projected an independent day 25.5 organoid dataset [59], generated using an adapted Multi-Factor protocol with PD173074- and SB431542-mediated inhibition of FGF receptor and TGFb/Activin/Nodal signaling, onto the 3D atlas using scArches (Fig. S5e). Mapping quality was consistently high across major cell types (Fig. 5i), enabling direct transfer of both broad and fine-grained annotations using CellTypist models trained on the atlas. This approach rapidly identified rare populations including WT1+BNC1+TBX18+ epicardial cells, which represent the organoid cell state with the highest similarity to fetal epicardium, as well as SHOX2+HCN4+ sinoatrial node-like pacemaker cells (Fig. 5j,k). Importantly, these annotations immediately enable downstream biological analyses. For example, the pacemaker cell population could be placed within the broader developmental context of the atlas and compared against equivalent populations across differentiation protocols, while the WT1+BNC1+TBX18+ population could be benchmarked against fetal epicardial reference states. Beyond cell-type identification, we leveraged the atlas to directly benchmark the maturity of this query dataset. Strikingly, despite being collected at day 25.5, the cardiomyocytes of the query organoid showed electrophysiological and calcium handling maturity comparable to cardiomyocytes of organoids from much earlier developmental stages (days 10–15) (Fig. 5l), while metabolic and structural maturity of the cardiomyocytes remained markedly limited (Fig. S5f). These results illustrate how atlas-guided benchmarking can reveal developmental differences that are not apparent from culture duration alone.

Together, these analyses demonstrate the versatility of the HHOCA as a resource for the field. In combination with data projection and automated label transfer, the atlas enables robust annotation of both common and rare cardiac cell states, quantitative benchmarking of developmental maturity, and systematic comparison across distinct cardiac differentiation strategies. More broadly, comparison across model systems revealed that micro-tissues favor metabolic maturation, organoids promote developmental, structural, and electrophysiological maturation through self-organized multicellular development, and embryoid bodies retain broader developmental potential before converging toward cardiac cell states. This highlights how a shared reference framework can be used to evaluate and compare increasingly diverse models of human cardiac development. To facilitate future studies, the custom CellTypist classifiers trained on the HHOCA are provided as a community resource, alongside the atlas itself, which is available via CELLxGENE [63] for interactive exploration. The atlas is additionally available via ArchMap [64] for reference-based mapping of new datasets (see Resource Availability).

### Inotropic signaling pathways remain underrepresented across cardiac in vitro models

Although cardiac organoids are increasingly used for drug discovery and safety assessment [65], our analyses indicate that they incompletely recapitulate adult cardiomyocyte states. We therefore asked whether this developmental immaturity affects their suitability for modeling pharmacological responses relevant to adult cardiovascular disease. As responsiveness to a given therapy necessarily depends on expression of its molecular targets, we evaluated the extent to which organoid-derived cardiomyocytes reproduce the drug-target repertoire of adult human cardiomyocytes. In addition to organoids generated using different protocols, we included cardiac micro-tissues because of their comparatively higher metabolic maturity.

Using the drug2cell pipeline [2], we compared predicted drug-target activity between adult cardiomyocytes and cardiomyocytes from organoids and micro-tissues based on the shared gene set. Because adult cardiomyocytes are typically profiled by single nucleus RNAseq whereas organoids and micro-tissues are profiled as whole cells, direct comparisons are confounded by the known bias of nuclear RNA sequencing towards long transcripts [66]. To minimize this effect, we quantified target expression relative to non-cardiomyocytes within each dataset using a cardiomyocyte-specificity (CM-specificity) score, which reduced but did not completely eliminate transcript-length bias (Spearman ρ = −0.52 versus −0.75; Fig. S6a,b,c). Of 80 drugs whose targets were enriched in adult cardiomyocytes, 46 showed significantly lower CM-specificity in organoid cardiomyocytes, and none showed higher specificity (FDR < 0.05) (Fig. 6a). Thus, although organoids recapitulate part of the adult cardiomyocyte drug-target repertoire (Fig. S6d), many clinically relevant targets remain incompletely represented.

**Figure 6:**
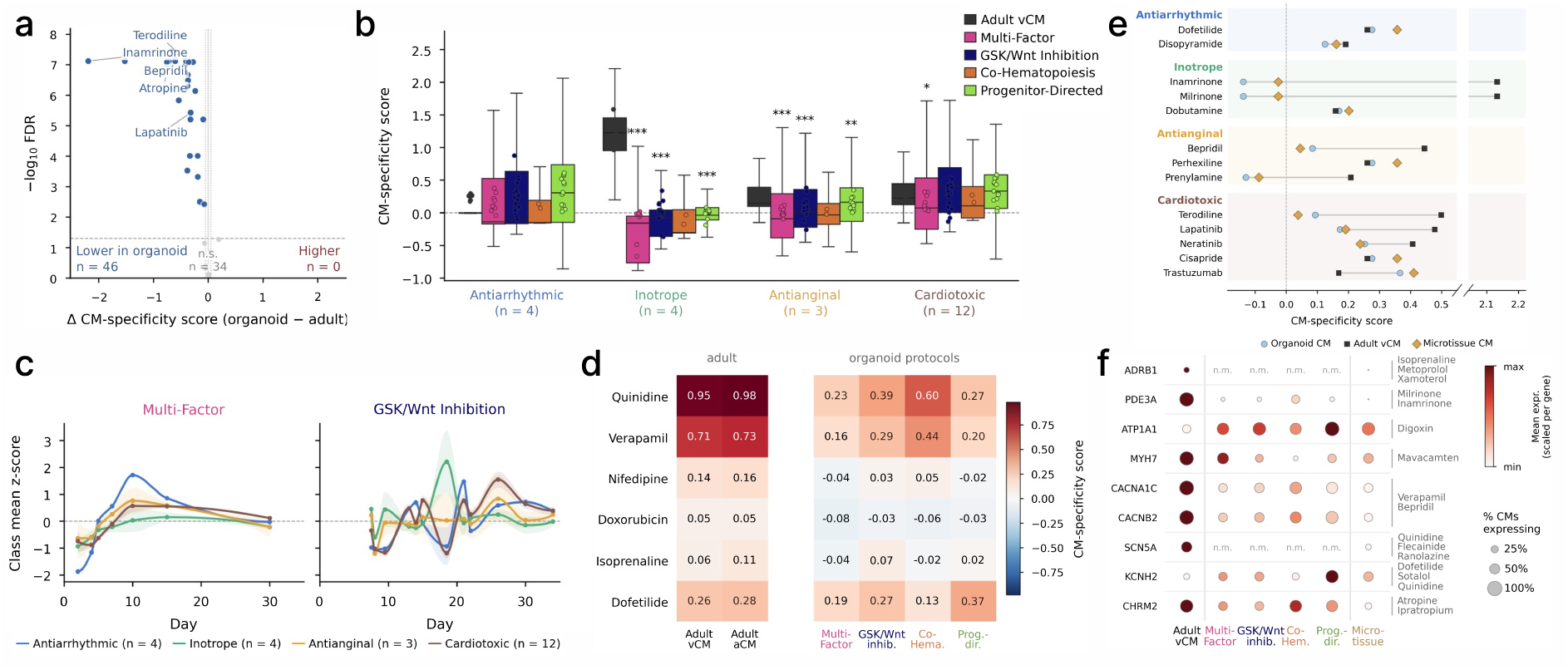
In-vitro cardiomyocytes capture antiarrhythmic and cardiotoxic drug targets but show a durable gap in inotropic signaling a, Organoid vs adult CMs (ventricular + atrial) for 80 drugs with adult-CM-enriched targets (of 2,447 ChEMBLdrugs in drug2cell package; adult CM score above the median and > 1.5× every non-CM adult cell type). Δ CM-specificity score (organoid − adult) vs −log10 FDR; two-sided Mann–Whitney U test on sample/donor means, Benjamini–Hochberg correction. **b,** CM-specificity score per drug class (classes with ≥ 3 drugs) per protocol; dots are samples/donors. Brackets: each protocol vs adult vCM (Mann–Whitney on sample means, BH-FDR; n/a, minimum reachable p > 0.05). **c,** Class mean of per-drug z-scored drug2cell scores over differentiation days in organoid CMs; bands, s.e.m. across drugs. **d,** CM-specificity score of drugs previously tested in heart organoids**. e,** Per drug: organoid CM, adult vCM and micro-tissue CM; arrow, organoid → adult; right, whether micro-tissue CMs are closer to adult vCM or to organoid CM. **f,** Expression of mechanism-of-action target genes in CMs: dot size, % expressing cells; color, mean log-normalized expression scaled per gene; n.m., gene not measured in that dataset. Adult = nuclei; % detection of long genes is higher in nuclei.

To identify which pharmacological pathways are most faithfully captured, we grouped cardioactive drugs into four categories: antiarrhythmics, inotropes, antianginals, and cardiotoxic compounds. The largest differences between adult and organoid cardiomyocytes were observed for inotropic targets. In adult cells, these targets showed the highest predicted activity of the four categories, whereas activity was markedly lower in all organoid protocols (all testable comparisons, p < 0.001; Fig. 6b). Antianginal target activity was likewise reduced in Multi-Factor and GiWi organoids (p < 0.001), but approached adult levels in Progenitor-Directed organoids. In contrast, antiarrhythmic targets were generally well represented, with Multi-Factor and GiWi organoids exhibiting activity comparable to adult cardiomyocytes and Progenitor-Directed organoids showing even higher activity. Cardiotoxic drug targets were largely similar between systems, although Multi-Factor organoids showed modestly reduced activity to adult cardiomyocytes (p < 0.05; Fig. 6b). Drug-specific effects within each category are shown in Fig. S6e.

Prolonged culture did not substantially improve the deficit in inotropic signaling. In Multi-Factor organoids, activity of most drug categories rose between day 2 and days 7–10 before plateauing. GiWi organoids showed a transient increase in inotropic target activity around day 18.5, followed by a subsequent decline (Fig. 6c), suggesting that time alone is insufficient to achieve adult-like representation of these pathways.

We next examined several cardioactive compounds that have been previously tested in cardiac organoids (Fig. 6d). Targets of the antiarrhythmics quinidine [67] and verapamil [68] displayed markedly greater CM-specificity in adult cardiomyocytes than in organoids (0.95 and 0.71 vs 0.16-0.60), whereas nifedipine [69] targets showed only minimal CM-specificity in organoids (−0.04 to 0.05 vs 0.14 in adult). In contrast, the hERG target of dofetilide displayed comparable CM-specificity in organoid (0.13 - 0.37) and adult cardiomyocytes (0.26), reaching the highest value in Progenitor-Directed organoids. Doxorubicin [65] targets exhibited low CM-specificity across all datasets, while assessment of isoprenaline responsiveness [69] was limited by the absence of *ADRB1* detection in the organoid datasets.

To determine whether enhanced metabolic maturity could compensate for these differences, we performed the same analysis in cardiac micro-tissue. Despite their increased metabolic maturity, micro-tissue cardiomyocytes did not display greater inotrope-target activity than organoid cardiomyocytes (Fig. 6e). Instead, CM-specificity scores for individual drugs closely resembled those observed in organoid cardiomyocytes, including the *PDE3* inhibitors milrinone and inamrinone (Fig. 6e).

Inspection of individual target genes revealed the molecular basis of these differences. *PDE3A*, a key mediator of inotropic signaling, was expressed in 93% of adult cardiomyocytes but only in 8–33% of organoid and 1% of micro-tissue cardiomyocytes. In contrast, *KCNH2* (hERG) and *ATP1A1* (digoxin) were detected in a greater fraction of organoid and micro-tissue cardiomyocytes than adult cells (Fig. 6f).

Together, these analyses demonstrate that cardiac organoids recapitulate specific components of the adult cardiomyocyte drug-target landscape while incompletely capturing others. In particular, expression of several clinically important electrophysiological targets, including hERG, closely approximated or exceeded adult cardiomyocyte specificity, supporting the use of organoids for interrogating these pathways. In contrast, inotrope-associated signaling, especially along the PDE3 axis, remained underrepresented across organoids and micro-tissues and was not rescued by prolonged culture. This unresolved deficit highlights a key aspect of adult cardiomyocyte biology that remains insufficiently modeled in vitro, while simultaneously identifying a tractable avenue for protocol refinement, given its strong dependence on differentiation strategy.

## Discussion

Here, we present the HHOCA, to our knowledge the first integrated single-cell transcriptomic atlas of hPSC-derived heart organoids. It comprises 422,059 cells from 14 publicly available and 2 unpublished single-cell transcriptomic datasets from 3D organoids and 2D monolayers, spanning 28 time points. Harmonized integration and marker-based cell type annotation enabled the identification of rare and previously difficult-to-resolve cell populations, including compact ventricular cardiomyocytes, multiple epicardial subtypes, pacemaker-like cells, and off-target ectodermal and neural populations. Beyond providing a comprehensive cellular census, HHOCA enables systematic evaluation of developmental fidelity, maturation state, protocol-specific biases, and multicellular communication across distinct cardiac model systems.

By anchoring organoid data to fetal and adult heart references, we were able to quantify how well these in vitro cells resemble their in vivo equivalent. A consistent finding across analyses is that most organoid-derived cell lineages closely resemble those of the early fetal heart. Across protocols, cells matched predominantly to early developmental stages, exhibited limited metabolic maturation, and lacked many transcriptional hallmarks of mature cardiac function. Ventricular cardiomyocytes in particular showed reduced expression of genes involved in oxidative phosphorylation, sarcomeric structure, contractile function, and excitation-contraction coupling relative to fetal reference cardiomyocytes. Importantly, organoid cells exhibiting greater similarity to in vivo fetal populations also consistently displayed more advanced maturation states, suggesting that insufficient maturation, rather than incorrect specification, is the main factor limiting fidelity. This distinction is encouraging since it implies that many core developmental programs are already established in current organoid systems and may be improved through strategies that specifically target maturation rather than lineage specification. Separately, given that increased cell stress has been reported as a common feature of organoid cultures compared to their primary counterpart [38], we re-examined our analyses throughout this study for evidence of a cell stress signature. We did not observe an increased cell stress phenotype in organoids relative to in vivo references.

Several observations suggest that tissue organization contributes importantly to the maturation process. Compared with stage-matched monolayer cultures, organoid ventricular cardiomyocytes were enriched for contractile and conduction programs, whereas 2D cardiomyocytes retained more proliferative states and largely failed to map to adult cardiomyocytes. Multicellular interactions, mechanical cues and tissue architecture are likely contributors [42], in line with our prediction of increasing epicardial derivative-to-cardiomyocyte signaling during organoid development. The immaturity also has practical consequences. The predicted activity of inotropic targets, particularly along the cAMP-PDE3 axis, was strongly reduced in all organoid protocols and did not increase with culture time or in micro-tissues, suggesting that current models may underestimate inotropic drug responses. Established maturation strategies, such as electrical or mechanical conditioning, metabolic media and hormonal stimulation [42][70], could now be benchmarked quantitatively against the fetal and adult references using the atlas.

At the level of cellular subtypes, correspondence to the fetal heart was more variable and provided insights into which developmental programs are successfully recapitulated and which remain incomplete. Among epicardial populations, only *WT1+BNC1+TBX18+* cells closely align with fetal mesothelial epicardium. However, the broader epicardial compartment emerged as one of the most faithfully reconstructed developmental systems within organoids. Consistent with observations in vertebrate development and hPSC-derived epicardial cultures [71]-[74], organoids reproducibly generated multiple epicardial states spanning specification, epithelial maintenance, EMT, and lineage diversification. Our pseudotime and gene expression analyses suggested a continuum extending from mesothelial epicardium through EMT-associated intermediates towards fibroblast-like fates. In particular, *WT1+TCF21+* cells displayed transcriptional features consistent with fibroblast commitment, in agreement with studies demonstrating a prominent fibroblast fate for TCF21-expressing epicardial derivatives [74][75][76]. The presence of these intermediate populations suggests that organoids capture dynamic epicardial developmental trajectories that are often transient and therefore difficult to study in vivo.

A similar theme emerged within the endothelial compartment. The hierarchy of fetal correspondence mirrors coronary development: venous endothelial cells matched their fetal counterparts most closely, whereas many cells resembled capillary-like or transitional states, and arterial cells expressed specification markers without acquiring the complete fetal arterial transcriptome. In vivo, coronary arterial endothelial cells arise through progressive reprogramming and remodeling of an immature, initially venous-like coronary plexus; pre-arterial identity emerges before perfusion, whereas subsequent arterial maturation is shaped by coronary blood flow, shear stress, and local tissue-derived signals [77]. The absence of a perfused vascular lumen and physiological flow in static organoids may therefore constrain endothelial network remodeling and full arterial maturation, providing a rationale for vascularized, microfluidic, or otherwise perfused organoid systems [78].

For cardiomyocytes, a particularly notable finding was the reproducible emergence of compact ventricular cardiomyocytes across independent organoid and monolayer datasets. Compact ventricular cardiomyocytes exhibited among the strongest correspondence to fetal ventricular populations, suggesting that essential aspects of ventricular wall patterning can emerge intrinsically during in vitro differentiation. The robust recovery of this population across multiple studies further supports the existence of conserved developmental programs that operate independent of protocol-specific differences. In contrast, more advanced programs associated with metabolic specialization, contractile maturation, and adult cardiomyocyte identity remained incompletely established, reinforcing the conclusion that terminal maturation rather than ventricular specification represents the principal developmental bottleneck.

Beyond evaluating developmental fidelity, HHOCA enabled systematic exploration of protocol-dependent differences. Although differentiation stage and protocol category explained more variation than the stem cell line identity, several biologically meaningful trends emerged. Multi-Factor protocols generated cardiomyocyte-rich organoids with greater electrophysiological and structural maturity, whereas GiWi-based protocols produced more heterogeneous tissues containing larger endothelial, mesenchymal and epicardial compartments, but also increased off-target populations. Within GiWi protocols, BMP4 and Activin A supplementation was associated with increased epicardial, fibroblast and pacemaker cell formation, consistent with the context-dependent roles of BMP/TGF-β signaling in early cardiac mesoderm patterning, proepicardial-epicardial specification, and sinoatrial-node differentiation [79][80][81]. Collectively, these findings reveal an important trade-off between cellular diversity and cardiomyocyte maturation that may guide protocol selection according to the intended biological application.

Several limitations should be considered. Although HHOCA represents the largest integrated collection of cardiac organoid single-cell data assembled to date, some protocol comparisons remain constrained by limited dataset numbers and unavoidable confounding variables. Differences in stem-cell lines, culture conditions, sequencing depth, and experimental design cannot be completely separated from protocol-specific effects. Furthermore, transcriptomic similarity does not necessarily imply functional equivalence, and future integration of spatial, epigenomic, proteomic, electrophysiological, and metabolic datasets will be essential to provide a more comprehensive assessment of model fidelity.

Finally, we demonstrate that HHOCA functions not only as a reference atlas but also as a quantitative benchmarking platform. Projection of engineered micro-tissues and embryoid bodies revealed how tissue assembly strategies shape lineage commitment and maturation, and the same approach can be applied to newly developed protocols, disease models and patient-specific cell lines. To this end, projection of a newly generated organoid dataset into the HHOCA reference space enabled rapid objective assessment of lineage composition, developmental state, maturation status, and cellular diversity. More broadly, HHOCA shifts the evaluation of cardiac organoids from qualitative descriptions of cell-type composition toward quantitative measurement of developmental fidelity and multicellular organization. As additional single-cell and spatial datasets become available [18][82], the atlas can be expanded to encompass later developmental stages, vascularized systems and immune-containing cardiac models. By providing a unified cell-state map of human cardiac development in vitro, HHOCA establishes a foundation for the standardized evaluation, comparison and rational engineering of next-generation human heart models.

## Material & Methods

### Data Collection and Harmonization

The atlas includes scRNA-Seq datasets from 14 publications [12]-[25] and two unpublished sources (Zengerle_X, Meier_X), comprising 13 three-dimensional (3D) datasets and 5 two-dimensional (2D) datasets. Two publications contributed both 3D and 2D samples. ScRNA-seq count matrices and associated metadata were obtained from the locations specified in the respective data availability statements, or directly from the authors in the case of unpublished data.

### Generation of the Zengerle_X organoid dataset

All human pluripotent stem cells (hPSCs) used in this study were cultured on Geltrex-coated (Gibco, A14133-02) tissue culture plates according to the manufacturer’s protocol. The hPSC line MRIi003-A was used for organoid generation. Cells were maintained under standard culture conditions (37°C, 5% CO₂) in Essential 8 medium (Gibco, A1517001) supplemented with 0.5% Penicillin/Streptomycin (Gibco, 15140-122). The culture medium was changed daily. Cells were passaged every four days using 0.5 mM EDTA (Invitrogen, AM92606) diluted in calcium- and magnesium-free PBS (PBS⁻/⁻; Gibco, 10010023). To enhance cell survival following passaging, 2 µM Thiazovivin (Sigma-Aldrich, SML1045) was added to the culture medium for 24 h.

Epicardioids were generated as previously described by Meier et al. [17], with the exception that VEGF (R&D, 293-VE-010) was omitted from the differentiation medium. Samples were collected on day 15 by pooling multiple organoids to obtain sufficient cell numbers for downstream analyses. For sample collection, single-cell dissociation of organoids were dissociated with papain as described previously [83]. The number of organoids and papain incubation time was adapted to the developmental age of organoids. Briefly, a 2× papain solution containing 40 U ml⁻¹ papain (Worthington Biochemical, LS003124) and 2 mM L-cysteine (Sigma-Aldrich, C6852) in PBS⁻/⁻ was activated for 10 min at 37°C and subsequently diluted 1:2 with PBS⁻/⁻ to generate the working solution. Spheroids were retrieved from the collagen gel and washed twice with 2 mM EDTA in PBS⁻/⁻. Dissociation was carried out in 750 µl papain working solution at 37°C with agitation at 750 rpm using a ThermoMixer (Eppendorf). Enzymatic activity was terminated by adding an equal volume of stop solution consisting of 1 mg ml⁻¹ trypsin inhibitor (Sigma-Aldrich, T9253) in PBS⁻/⁻. Samples were mechanically dissociated by repeated pipetting to obtain a single-cell suspension, filtered through a 40-µm cell strainer, and washed with 5 ml of 1% BSA (Gibco, 15260037) in PBS⁻/⁻. Following centrifugation at 200 × g for 3 min, cells were resuspended in 100 µl of 0.5% BSA in PBS⁻/⁻ and counted using trypan blue.

Subsequently, single-cell RNA sequencing libraries were prepared with a targeted cell recovery of 6000 cells using the Chromium Next GEM Single Cell 3′ Kit v3.1 (10x Genomics) according to the manufacturer’s instructions. cDNA quality was assessed using a Bioanalyzer (Agilent) with the High Sensitivity DNA Kit (Agilent, 5067-4626). Libraries were quantified using the KAPA Library Quantification Kit (KAPA Biosystems, KK4824) according to the manufacturer’s protocol. Following quantification, libraries were pooled and sequenced on an Illumina NovaSeq SP flow cell using 150-bp paired-end sequencing, with 28 cycles for Read 1, 91 cycles for Read 2, 8 cycles for the i7 index read, and no i5 index read. Sequencing was performed to a depth of ca. 33,000 reads per cell.

### Generation of the Meier_X organoid dataset

Epicardioid differentiation has been performed according to the protocol described by Meier et al. [17], without VEGF addition. The hPSC line MRIi003-A was used for organoid generation. Organoids were dissociated into single cells as described above and single cells were resuspended in 500 µl of 0.5% BSA in PBS⁻/⁻ before counting with trypan blue.

Single-cell RNA sequencing libraries were prepared with a targeted cell recovery of 8000 cells using the Chromium Next GEM Single Cell 3′ Kit v3.1 (10x Genomics) according to the manufacturer’s instructions. cDNA quality was assessed using a Bioanalyzer (Agilent) with the High Sensitivity DNA Kit (Agilent, 5067-4626). Libraries were quantified using the KAPA Library Quantification Kit (KAPA Biosystems, KK4824) according to the manufacturer’s protocol. Following quantification, Libraries were pooled and sequenced using a NovaSeq S1 flow cell (Illumina) with 150-base pair (bp) paired-end reads with 28 cycles for read 1, 91 cycles for read 2, 8 cycles for i7 and 0 cycles for i5 and with a read depth of at least 25,000-30,000 paired-end reads per cell.

### Data Preprocessing

Unless otherwise stated, all data processing and analyses were performed using Scanpy (v1.11.5) [84]. For quality control and filtering, the number of genes expressed and the number of counts per cell were used as metrics. Doublet detection was carried out with Scrublet [85], run separately for each dataset with the sample specified as the batch key. Cells predicted as doublets and cells with fewer than 100 genes expressed or fewer than 750 total counts, were removed. After filtering, all datasets were merged into a common feature space, of which only ∼11,000 genes were present across all datasets. This reduction was primarily driven by the datasets of Schmidt et al. and Feng et al., which had a limited feature space of approximately 11,000 genes. For this reason, we additionally created a second version of the atlas excluding these two datasets, allowing for a larger gene space. In total, the resulting atlas comprises 422,059 cells, of which 316,711 originate from 3D organoids and 105,348 from 2D monolayers. Raw counts were then normalized such that each cell’s total count matched the median total count across all cells prior to normalization, and were subsequently log-transformed.

### Data Integration Benchmark

Data integration was performed to generate three separate embeddings: one combining 3D and 2D data, and one each for 3D and 2D data separately. Four integration methods - scVI [86], scANVI [87], scPoli [88], and Harmony [89] -were benchmarked using the Single Cell Atlassing Toolbox (scAtlasTb) [40]. Each method was run using three different highly variable gene (HVG) selection flavors with 3,000 genes being selected: seurat, seurat_v3, and seurat_v3_paper, with the batch key for HVG selection set to sample in each case. For the integration of 3D and 2D, the HVGs were computed separately, and the intersection of the two HVG selections was taken as the set of HVGs. For every combination, two batch variables were additionally tested for the integration methods themselves: dataset and sample. scVI, scPoli, and Harmony were run in an unsupervised manner, whereas scANVI incorporated the broad level 1 annotation as a label key. For scVI and scANVI, additional hyperparameters were benchmarked, including the number of epochs and latent space size. The best-performing integration method was determined based on 13 scIB metrics [41]. scVI outperformed the other methods for both the combined 3D+2D (n_latent=20) and the 2D-only (n_latent=10) embeddings, with seurat_v3_paper yielding the best-performing HVG selection for 3D+2D and seurat_v3 for 2D-only. For the 3D embedding, scANVI outperformed the other methods, using the seurat_v3_paper HVG selection with n_latent=20. Across all best-performing integrations, sample rather than dataset was used as the batch key.

### Cell-Type Annotation

A marker gene list was compiled from the 14 publications from which the data were obtained (Table S2). Genes were categorized into three groups: consensus markers, mentioned for a single cell type across more than one publication; publication-specific markers, used in only one publication; and ambiguous markers, used to define multiple cell types across different publications. This marker gene list was used to perform the cell type annotation of all three levels. Level 2 and level 3 annotations were performed separately for 2D and 3D data, using their respective integrated embeddings. Annotation was based on Leiden clustering combined with differential expression analysis (Wilcoxon rank-sum test) and gene scores. For the level 3 annotation, cell states were defined only for cardiomyocytes, endothelial cells, mesenchymal cells, and epicardial cells in the 3D data. In the 2D data, due to lower cellular heterogeneity, level 3 annotation was restricted to cardiomyocytes.

### Compositional analysis

To analyse changes in cell-type composition across developmental stages, we calculated the proportion of cells in each cell type (level-2 annotation) for every dataset and stage, separately for 2D and 3D cultures. Samples with fewer than 10 cells were excluded. Zero counts were replaced using the geometric Bayesian-multiplicative method (zCompositions v1.5.0.5) [90], and proportions were CLR-transformed (compositions v2.0.9) [91]. For each cell type, we compared CLR-transformed values for the early and middle stages with the late stage using linear models. P-values were corrected for multiple testing with the Benjamini-Hochberg method, and changes with FDR < 0.05 were considered significant. Effect sizes are reported as log2 fold changes in mean proportion relative to the late stage.

### Differential gene expression analysis for Cardiac Subtype Diversity Analysis

Differential expression was tested at the pseudobulk level between 2D and 3D models in ventricular cardiomyocytes, separately for the middle and late stages and between developmental stages in 3D cardiomyocytes (early and late, each compared with middle). For each group, raw counts were summed across all cells from the same dataset and differentiation day to give one pseudobulk sample. Only samples with at least five cells were included, and each group needed at least three samples. Genes were kept if their mean log-normalized expression was at least 0.0125 in at least one of the two groups (counts normalized to 10,000 per cell, log1p-transformed, and averaged across samples). Analyses were performed in R (version 4.5.1) with the edgeR package (version 4.6.3) [92]. Library sizes were normalized with the trimmed mean of M-values (TMM) method [93]. Counts were modeled with a negative binomial generalized linear model, with dispersions estimated separately for each comparison, and differences were tested with quasi-likelihood F-tests [94]. P-values were corrected for multiple testing with the Benjamini-Hochberg method. Genes were classified as differentially expressed if the false discovery rate (FDR) was ≤ 0.05 and the absolute log₂ fold change was ≥ 0.75.

### Gene Ontology enrichment for Cardiac Subtype Diversity Analysis

#### 2D versus 3D comparison

Functional enrichment was assessed by preranked Gene Set Enrichment Analysis (GSEA) using gseapy (v1.1.11, Python 3.10) [95]. For each stage (middle and late), genes from the edgeR comparison of 3D versus 2D were ranked by sign(log2FC) × −log10(p-value), without a fold-change cutoff. Ensembl-only identifiers, mitochondrial, LINC and ribosomal protein genes were excluded, and duplicate gene symbols were collapsed to the entry with the largest absolute log2FC. GSEA was performed against GO Biological Process gene sets (GO_Biological_Process_2026), restricted to 10-500 genes, using 1,000 permutations. Positive NES indicated enrichment in 3D and negative NES enrichment in 2D. Terms with FDR ≤ 0.05 were considered significant and classified as middle-, late-stage or shared according to the stage(s) in which they were significant. For visualization, representative terms were selected.

#### program timing in 3D cardiomyocytes

Changes in functional programs were assessed using edgeR comparisons between stages (middle vs early, late vs middle and late vs early) in 3D cardiomyocytes. Comparisons were oriented such that positive log2FC indicated higher expression in the later stage. Genes were ranked and filtered as described above, and preranked GSEA was performed against GO Biological Process, Molecular Function and Cellular Component gene sets (2026 releases), using gene sets of 10-500 genes, 1,000 permutations and seed 0. Terms with FDR q ≤ 0.05 were considered significant. Terms significant in multiple comparisons were retained once, prioritising late vs early, followed by late vs middle and middle vs early, to define a single leading-edge gene set. Biological Process terms representing proliferation, metabolism, specification, conduction and contractile maturation were selected for display. Terms that did not reach significance in the selected comparison are reported with their best comparison and FDR (Table S4).

For each selected term, program activity was scored per cell using the leading-edge genes with scanpy.tl.score_genes, which subtracts the mean expression of an expression-matched control gene set from log-normalized counts. Scores were averaged per sample, with each sample defined as one dataset at one differentiation day containing at least 20 cardiomyocytes. Sample scores were centered across scored programs and then centered within the dataset to reduce between-dataset effects. Datasets represented by only one day window were excluded from the within-dataset centering. Samples were grouped into three differentiation windows (days 0-10, 11-20, 21-35); windows containing fewer than two samples or 500 cells were excluded. Scores were subsequently z-scored across windows for each program.Because the scored genes were derived from the same stage comparisons, program profiles were interpreted primarily in terms of their relative timing and ordering rather than effect magnitude.

### Multinichenet

Cell-cell interactions between conditions were inferred using MultiNicheNet (multinichenetr v2.1.0, R v4.5.1) [43]. The input was a SingleCellExperiment [96] of raw counts from 3D cardiac organoids generated with Multi-Factor or GSK/Wnt-inhibition protocols (29 samples; 9 early, 10 middle,10 late). Analysis was restricted to 24 cardiomyocyte and epicardial cell states (cell_type_level3), each considered as both sender and receiver. Cell types were retained in a sample only if they were represented by at least 10 cells, and only cell types detected in at least three samples per group were included in the analysis. Genes were considered expressed in a given cell type if detected in at least 0.05 of its cells in at least 0.5 of samples within a stage. Prior knowledge on intercellular signaling was obtained from the NicheNet v2 model [97]: the human ligand-receptor network (lr_network_human_allInfo_30112033.rds, Zenodo record 10229222), reduced to unique ligand-receptor pairs, and the ligand-target regulatory potential matrix (ligand_target_matrix_nsga2r_final.rds, Zenodo record 7074291). Ligand and receptor gene aliases were harmonized to current official gene symbols using convert_alias_to_symbols from nichenetr prior to analysis. Differential expression between conditions was computed on pseudobulk profiles using edgeR within the MultiNicheNet framework, with contrasts [’early-late’, ‘middle-early’,’late-early’, ‘middle-late’, ‘late-middle’] and with protocol type as a batch covariate.

Ligand activity was predicted for each receiver cell type from its differentially expressed genes, using thresholds of logFC ≥ 0.5 and adjusted_p ≤ 0.05, using the top 250 predicted targets per ligand, separately for up- and down-regulated genes. Interactions were ranked with the default (“regular”) MultiNicheNet prioritization, which combines differential ligand and receptor expression, scaled ligand activity, cell-type specificity and the fraction of samples expressing both partners.

For each stage, the interaction score was defined as the highest prioritization score across the two contrasts at that stage. An interaction was considered prioritized if its score was ≥ 0.80 and both ligand and receptor were expressed in ≥ 50% of samples. Cell-cell communication was summarised as the number of prioritized interactions for each sender-receiver pair and visualized as chord diagrams, with separate diagrams for each stage and a pooled analysis. In the pooled analysis, interactions prioritized at multiple stages were counted once. Only interactions between distinct cell types were included, and sender-receiver pairs contributing < 1% of the total interactions in a given panel were omitted from visualization.

For stage-resolved dot plots, each interaction was assigned to the stage at which it reached its highest prioritization score (peak stage), and the three highest-ranked interactions for each peak stage were shown across all three stages. Dot color indicates the prioritization score and dot size the fraction of samples expressing both ligand and receptor. Additional dot plots summarised interactions involving vCMs, separated into interactions sent by, received by, or exchanged with vCMs.

For the summary figure, 15 vCM-associated interactions (five per stage) were selected from the prioritized set to represent signaling pathways with established or proposed roles in cardiac development. Predicted downstream effects were assessed using NicheNet regulatory potential scores between each ligand and target genes upregulated in the corresponding receiver cell type. The top five targets per ligand were shown for ligands received by vCMs, and the top six targets per ligand and receiver were shown for ligands sent by vCMs. For each ligand-target pair, the highest regulatory potential score across the contrasts within the corresponding stage was used.

### Cross-dataset neighborhood similarity analysis

Organoid cells were compared to fetal and adult heart atlases using a neighborhood comparison approach implemented in scrabbitr (v0.1.0) [50], based on miloR neighborhoods [56]. Because all datasets were human, no ortholog mapping was required, and analyses were restricted to genes shared between datasets. Raw counts were log-normalized with scuttle (logNormCounts). For each dataset, a k-nearest neighbour (kNN) graph (k = 35) was built on the scVI latent space (first 30 dimensions, or 20 where the embedding had fewer dimensions). neighborhoods were defined with makeNhoods by randomly sampling 5% of cells as index cells (refined = FALSE) and assigning each index cell’s 35 nearest neighbours to its neighborhood. Highly variable genes (top 5,000 per dataset) were selected with scrabbitr’s selectNhoodFeatures, and the intersection between organoid and atlas was retained. For each neighborhood, mean log-normalized expression was calculated and converted to gene-specificity scores by dividing each gene’s neighborhood mean by its mean across all neighborhoods in that dataset. Pearson correlations between the gene-specificity profiles of all organoid and atlas neighborhoods gave a neighborhood similarity matrix. Each neighborhood was then assigned its most highly correlated neighborhood in the other dataset, in both directions. For visualization, neighborhoods were plotted at the UMAP coordinates of their index cells in each dataset and colored by the most frequent cell-type annotation among their member cells (organoid: level-1 annotation; fetal atlas: mid-grain annotation). Each neighborhood’s best-matching neighborhood in the other dataset was indicated by a connecting line (scrabbitr plotTrajMappings). The fetal embedding was reflected along the y-axis and offset for display only, and cell types with the same label in both datasets were given the same color.

### Maturity Score

To compute the maturity score, the pyUCell package (v0.7.3) [51] was used. The pyUCell package provides rank-based gene-signature scoring using the Mann-Whitney U statistic. The pyUCell score allows incorporating positive and negative gene sets and weighting the contribution of the negative set with the weighting parameter w_neg. For computing the maturity scores, the parameter missing_genes was set to “skip” and w_neg was set to 1. max_rank was defined for each dataset based on its median n_genes_by_counts. Genes used to compute maturity scores were curated from the literature (Table S3) [42][98]-[104].

### SysVI Integration and Presence Score

We used SysVI (scvi-tools v1.4.2) [52][105] to integrate reference (organoid/monolayer) and query (fetal/adult) datasets. The model was run on shared HVGs. HVGs were calculated separately for reference and query with batch_key set to the sample. The intersection of both HVG sets was then used for SysVI integration. The batch_key in SysVI.setup_anndata was set to the reference/query, respectively, while categorical_covariate_keys was set to the sample. The SysVI model was run with embed_categorical_covariates set to True. kl_weight and cycle_weight was set to 5 for organoid integrations, while kl_weight was set to 10 and cycle_weight to 2 for monolayer integrations. The presence score was computed for each cell in the reference as previously described [38].

### Differential Gene Expression between Fetal and Organoid

Differential Gene Expression analysis was performed between organoid and fetal groups, stratified by organoid similarity score (threshold: 0.5). More similar and less similar organoids were also compared directly to one another. DEG was performed using a Wilcoxon Rank Sum test with scanpy.tl.rank_genes_groups.

### Multi Omics data analysis-MOFA

For the integrative analysis, we restricted the data to 3D organoid samples generated with GSK/Wnt inhibition (GiWi; n = 23) or Multi-Factor (n = 10) protocols, excluding samples from the Progenitor-Directed and Co-Hematopoiesis protocols. Each cell type was modeled as its own feature set.To capture variation below the main annotation, we added each sample’s level-2 cell-type composition to the corresponding cell type after a centered log-ratio (CLR) transformation.

MOFA requires one expression profile per sample and cell type [55]. We therefore summed gene counts across cells of the same type within each sample and normalized the resulting profiles to the same total count to account for differences in sequencing depth.

To reduce sparsity, we excluded epithelial and hematopoietic cells, which were too rare to model samples. For the remaining cell types, we retained genes that satisfied at least one of two detection criteria:

● detected in more than 50% of the cells of that type and in more than 330 cells in total, or
● detected in more than 40% of the cells of that type and in more than 660 cells in total.

These cut-offs balance two goals. Each retained gene is robustly detected across many cells and samples, and every cell type still keeps enough genes for downstream modeling.

We log-transformed and quantile-normalized the filtered counts so expression distributions were comparable across samples. In total, we kept 13,298 genes across all cell types. These made up the cell-type-specific single-cell views in the MOFA model.

Cell-type composition was expressed as centered log-ratio (CLR)-transformed values for each sample, which were used directly in subsequent analyses without further transformation.

After preprocessing, the dataset contained 13,340 features across 8 views: 7 cell-type views from the scRNA-seq data (13,298 genes in total) and one view of 42 cell-state composition features. We then quantile-normalized all views feature-wise, mapping each feature’s values to the quantiles of a standard normal distribution.

We fitted the MOFA model using the R/Bioconductor package MOFA2 (version 1.2.2) [55], with 10 factors and a maximum of 50,000 iterations, keeping other parameters at their default settings. We selected 10 factors because they collectively explained more than 80% of the total variance across feature views.

Pathway enrichment was performed on the MOFA feature weights of factors 1-5 using `run_enrichment` from MOFA2 (version 1.2.2) and gene sets from the Enrichr Reactome_2023 Pathways, with less than 20% of their genes in the feature set were excluded. Pathways related to cardiomyocyte development were selected for visualization.

### Protocol Analysis

Differential abundance testing was performed using Milo [56], as implemented in pertpy (v1.0.3) [106]. Cell types with fewer than 10 cells in a subset were removed prior to testing. The neighborhood graph was built on the integrated embeddings, with neighborhoods defined by sampling 10% of the cells as index cells. Each neighborhood was annotated by its majority cell type. Neighborhoods in which the majority label accounted for less than 60% of cells were labeled “Mixed” and excluded from further analysis. The edgeR solver was used for differential abundance testing. Neighborhoods were called differentially abundant at a spatial FDR < 0.05. Code for the Milo differential abundance analysis and for generating figures was developed with the assistance of Claude Sonnet 5 (Anthropic), and was reviewed and validated by the authors.

Differential Gene Expression analysis was performed using Wilcoxon Rank Sum tests. The resulting differential expression results were then used for gene set enrichment analysis (GSEA), performed using the gseGO function from the R package clusterProfiler (v4.18.4) [107], with minGSSize set to 3, pvalueCutoff set to 0.05, and pAdjustMethod set to “BH”.

### Evaluating Diverse Differentiation Strategies

To evaluate diverse differentiation strategies using the HHOCA as a reference, we included a micro-tissue dataset [25], an embryoid body dataset [20], and an independent organoid dataset [59]. In addition, we comprehensively reviewed whether suitable disease or perturbation cardiac organoid datasets were publicly available for potential mapping onto the atlas. Following careful evaluation, we concluded that currently available datasets did not constitute validated disease or perturbation models appropriate for benchmarking against the HHOCA, and were therefore excluded from this study.

The scRNA-seq count matrices and associated metadata of the data used for data projection were obtained from the locations specified in the respective data availability statements. For quality control and filtering, cells predicted as doublets via Scrublet and cells with fewer than 100 genes expressed or fewer than 750 total counts, were removed.

Reference mapping onto the organoid atlas was performed using scArches (scvi-tools v1.4.2) [60][105], using the scANVI model from the 3D integration. Label transfer of level 1 cell types was conducted using CellTypist (v1.7.1) [61] with a custom CellTypist model trained on the organoid atlas. The custom model was trained on data that was normalized to 10,000 counts per cell and log1p-transformed. Label transfer was then performed using celltypist.annotate, with majority_voting set to True, mode set to “prob match”, and both p_thres and min_prob set to 0.4. To evaluate the quality of the query-to-reference mapping, MapQC (v0.1.1) [62] scores were calculated using mapqc.run_mapqc, with n_nhoods set to 2000, k_min set to 50, and k_max set to 1000.

For the Differential Gene Expression analysis between micro-tissue and organoid data, the organoid atlas was subset to organoids with day 30 or older to ensure comparability. Differential Gene Expression analysis was performed using Wilcoxon Rank Sum tests. The resulting differential expression results were then used for gene set enrichment analysis (GSEA), performed using the gseGO function from the R package clusterProfiler (v4.18.4) [107], with minGSSize set to 3, pvalueCutoff set to 0.05, and pAdjustMethod set to “BH”. Gene scores of neural, hepatic, and cardiac fate were computed using scanpy.

### Drug target scoring

Drug-target activity was scored per cell with drug2cell (v0.1.2) [2] using its built-in ChEMBL drug-target set. We scored the adult heart atlas [2], restricted to 10x 3′ v3 single-nucleus data (n = 190,497), the heart organoid atlas (n = 316,711), and cardiac micro-tissues (n = 7,658) [25]. Duplicate gene symbols were collapsed to the copy with the highest total counts. We retained only drugs scored on identical target genes across all three datasets (n = 1,811). Count matrices were normalized to 10⁴ counts per cell and log1p-transformed in scanpy (v1.11.5); the micro-tissue data were already log-normalized and used as provided. Scores were computed on the normalized matrix (use_raw=False). Drugs were called CM-enriched (n = 80) if their mean score in adult vCM or aCM was above the median across drugs and > 1.5-fold higher than in every non-CM cell type (≥ 50 nuclei). The CM-specificity score was defined per sample or donor as the mean CM score minus the mean non-CM score of the same sample or donor. To assess nuclear versus whole-cell bias, we used one adult donor (D11) [2] profiled by both snRNA-seq and scRNA-seq. Per-cell-type scores were compared between nuclei and cells, and per-drug differences were correlated between cell types (Spearman) (Fig. S6a, b). Transcript-length bias was assessed by Spearman correlation between organoid − adult differences and median target-gene length (GENCODE v44) (Fig. S6c). Agreement between organoid and adult vCM CM-specificity scores across all drugs was quantified by Pearson correlation (Fig. S6d). We compared organoid samples (n = 51) and adult donors (n = 7) using two-sided Mann–Whitney U tests on sample/donor means, with Benjamini–Hochberg correction (FDR < 0.05); Co-Hematopoiesis organoids (n = 2) were not testable. We grouped 23 drugs into four pharmacological classes (antiarrhythmics, inotropes, antianginals, cardiotoxic compounds), and analysed drugs previously tested in organoids separately (Fig. S6e). For time courses, we z-scored mean daily scores per drug and averaged them per class. Micro-tissues (one sample) were compared descriptively. The fraction of CMs expressing each target gene (counts > 0) was computed per dataset.

### Data visualization

Code used to generate figures was developed with the assistance of Claude Sonnet 5 (Anthropic), and was reviewed and validated by the authors.

## Supplementary Information

HHOCA Metadata Information (Table S1): *HHOCA_S1_Metadata.xlsx*

Table of genes used for annotation (Table S2): *HHOCA_S2_Marker_Gene_List.xlsx*

Table of genes used for maturity scores (Table S3): *HHOCA_S3_Maturity_Genes.xlsx*

DEG list 2D vs 3D organoids per stage, 3D CM and GSEA analysis (Table S4): *HHOCA_S4_DEG_GSEA.xlsx*

## Resource Availability

The atlas (3D+2D, 3D only, and 2D only) will be made available on CELLxGENE for interactive exploration. The 3D atlas, along with its reference model, will additionally be made available on ArchMap for reference-based mapping of new datasets. The CellTypist models for level 1, level 2, and level 3 annotation will be made publicly available at 10.5281/zenodo.23802040 upon publication.

## Acknowledgments/Funding

This work was supported by the European Research Council (ERC; grants: 788381/BIOCARD and 101141820/EPICURE to A.M., 101054957/DeepCell to F.J.T.), the European Union (EU Horizon; contracts 101080204/GEREMY to A.M., 101056712/HEAL, 101257413/HEARTCORE and 101288899/iNDUCARE to R.Z.), the German Research Foundation (DFG; Transregio Research Units 152 and 267 to A.M. and K.-L.L., Excellence Clusters 3092/BioSysteM and 3113/NUCLEATE to A.M., ZW 64/4-2 to R.Z.), Federal Ministry of Research, Technology and Space (BMFTR; grants: 01EK1601A, 031L0249, 01EK2108A and 03ZU2202HB/curATime to R.Z., 03ZU1201BH/CNATM to A.M.), the German Center for Cardiovascular Research (DZHK; grant FKZ 81Z0600601 to A.M. and K.-L.L., grants 81Z0600111 and 81Z0600112 to M.H and F.J.T.),Lower Saxony (‘Zukunft Niedersachsen’ grant: ZN4092, ‘Förderung aus EFRE-Mitteln’ grants: ZW3-87035144, ZW 7-87056735 to R.Z.), and NCR3s UK (grant: NC/Z500707/1 to R.Z.). L.D. received funding from the PREPARE Medical Scientist Program (funded by Hannover Medical School and the Ministry of Science and Culture of the State of Lower Saxony, MWK). S.O., V.D., K.T., M.G., and L.D.M. are supported by the Helmholtz Association under the joint research school “Munich School for Data Science - MUDS”.

## Author Contributions

S.O. and V.D. contributed equally. M.G., L.D.M., and S.O. compiled the list of datasets to integrate into the atlas. S.Z, A.B.M., and T.D. generated the additional heart organoid datasets. S.O., M.G., L.D.M., T.A., A.V.D.B., and S.M. performed data collection and preprocessing, supervised by L.H.; T.A., A.V.D.B., and S.M. were additionally supervised by K.T. and V.D. S.O. and A.V.D.B. performed integration of the atlas. S.O., S.M., L.D., S.Z. and J.G. performed cell type annotation. V.D. performed analysis of ventricular cardiomyocyte programs. S.O., V.D., and T.A. performed analysis of heart organoid fidelity. S.O., V.D., J.G., C.M., L.X., L.S., and J.L.B. performed protocol analysis, with J.G., C.M., L.X., and L.S. supervised by S.O. and J.L.B.. S.O. performed analysis of differentiation strategies using the atlas as a reference. V.D. and L.H. performed drug-target activity analysis. F.J.T, L.D. and K.T. edited the manuscript. S.O., V.D., L.H., A.M., and M.H. wrote the manuscript. F.J.T., A.M., M.H., R.Z., L.H., K.-L.L. conceived the study and supervised analysis.

## Declaration of Generative AI and AI-Assisted Technologies in the Manuscript Preparation Process

During the preparation of this work, the authors used Claude Sonnet 5 and Opus 5.5 (Anthropic) to assist in generating code for the differential abundance analysis in the protocol comparison section, and to assist in generating code used to create figures throughout the manuscript, as well as to improve code readability and consistency. After using this tool, the authors reviewed and edited the content as needed and take full responsibility for the content of the published article.

## References

1. Litviňuková, M. et al. Cells of the adult human heart. Nature 588, 466–472 (2020).

2. Kanemaru, K. et al. Spatially resolved multiomics of human cardiac niches. Nature 619, 801–810 (2023).

3. Timmis, A. et al. European Society of Cardiology: cardiovascular disease statistics 2025. Eur Heart J 47, 3499–3543 (2026).

4. Main causes of mortality: Health at a Glance 2025. OECD https://www.oecd.org/en/publications/health-at-a-glance-2025_8f9e3f98-en/full-report/main-causes-of-mortality_17b5a2df.html (2025).

5. Zhang, X. et al. Global, regional, and national burden of congenital heart disease, 1990-2021: a systematic analysis for the global burden of disease study 2021. Eur J Pediatr 184, 253 (2025).

6. Schulz-Menger, J. et al. 2025 ESC Guidelines for the management of myocarditis and pericarditis: Developed by the task force for the management of myocarditis and pericarditis of the European Society of Cardiology (ESC)Endorsed by the Association for European Paediatric and Congenital Cardiology (AEPC) and the European Association for Cardio-Thoracic Surgery (EACTS). European Heart Journal 46, 3952–4041 (2025).

7. Parnasa, E. et al. Reversing the pipeline: a ‘human-first’ multi-omics approach to cardiovascular discovery. Exp Mol Med 58, 2859–2872 (2026).

8. Milani-Nejad, N. & Janssen, P. M. L. Small and large animal models in cardiac contraction research: Advantages and disadvantages. Pharmacology & Therapeutics 141, 235–249 (2014).

9. Krishnan, A. et al. A detailed comparison of mouse and human cardiac development. Pediatr Res 76, 500–507 (2014).

10. Van Norman, G. A. Limitations of Animal Studies for Predicting Toxicity in Clinical Trials: Is it Time to Rethink Our Current Approach? JACC Basic Transl Sci 4, 845–854 (2019).

11. Atkins, J. T. et al. Pre-clinical animal models are poor predictors of human toxicities in phase 1 oncology clinical trials. Br J Cancer 123, 1496–1501 (2020).

12. Dardano, M. et al. Blood-generating heart-forming organoids recapitulate co-development of the human haematopoietic system and the embryonic heart. Nat Cell Biol 26, 1984–1996 (2024).

13. Drakhlis, L. et al. Human heart-forming organoids recapitulate early heart and foregut development. Nat Biotechnol 39, 737–746 (2021).

14. Galdos, F. X. et al. Combined lineage tracing and scRNA-seq reveals unexpected first heart field predominance of human iPSC differentiation. eLife 12, e80075 (2023).

15. Ho, B. X. et al. Robust generation of human-chambered cardiac organoids from pluripotent stem cells for improved modelling of cardiovascular diseases. Stem Cell Res Ther 13, 529 (2022).

16. Feng, W. et al. Computational profiling of hiPSC-derived heart organoids reveals chamber defects associated with NKX2-5 deficiency. Commun Biol 5, 399 (2022).

17. Meier, A. B. et al. Epicardioid single-cell genomics uncovers principles of human epicardium biology in heart development and disease. Nat Biotechnol 41, 1787–1800 (2023).

18. O’Hern, C. et al. Human heart-macrophage assembloids mimic immune-cardiac interactions and enable arrhythmia disease modeling. Cell Stem Cell 32, 1671–1690.e13 (2025).

19. Schmidt, C. et al. Multi-chamber cardioids unravel human heart development and cardiac defects. Cell 186, 5587–5605.e27 (2023).

20. Snabel, R. R. et al. Cardiac differentiation roadmap for analysis of plasticity and balanced lineage commitment. Stem Cell Reports 20, 102422 (2025).

21. Tian, Y. et al. Generation of mature epicardium derived from human-induced pluripotent stem cells via inhibition of mTOR signaling. Nat Commun 16, 5902 (2025).

22. Volmert, B. et al. A patterned human primitive heart organoid model generated by pluripotent stem cell self-organization. Nat Commun 14, 8245 (2023).

23. Wang, F., Zou, X., Zheng, H., Kong, T. & Pei, D. Human epicardial organoids from pluripotent stem cells resemble fetal stage with potential cardiomyocyte-transdifferentiation. Cell Biosci 15, 4 (2025).

24. Zawada, D. et al. Retinoic acid signaling modulation guides in vitro specification of human heart field-specific progenitor pools. Nat Commun 14, 1722 (2023).

25. Zhang, F. et al. Single-cell atlas of multilineage cardiac organoids derived from human induced pluripotent stem cells. life. Med. 1, 179–195 (2022).

26. Eschenhagen, T. et al. Three-dimensional reconstitution of embryonic cardiomyocytes in a collagen matrix: a new heart muscle model system. The FASEB Journal 11, 683–694 (1997).

27. Guo, N.-N. et al. Early prediction of the differentiation potential during the formation of human iPSC-derived embryoid bodies. Biochemical and Biophysical Research Communications 516, 673–679 (2019).

28. Itskovitz-Eldor, J. et al. Differentiation of Human Embryonic Stem Cells into Embryoid Bodies Comprising the Three Embryonic Germ Layers. Mol Med 6, 88–95 (2000).

29. Eschenhagen, T. et al. 3D engineered heart tissue for replacement therapy. Basic Res Cardiol 97 Suppl 1, I146–152 (2002).

30. Gokhan, I., Blum, T. S. & Campbell, S. G. Engineered heart tissue: Design considerations and the state of the art. Biophysics Rev. 5, 021308 (2024).

31. Brade, T., Pane, L. S., Moretti, A., Chien, K. R. & Laugwitz, K.-L. Embryonic Heart Progenitors and Cardiogenesis. Cold Spring Harb Perspect Med 3, a013847 (2013).

32. Mendjan, S., Deyett, A. & Yelon, D. Coordination of cardiogenesis in vivo and in vitro. Nat Rev Mol Cell Biol 27, 19–34 (2026).

33. Lewis-Israeli, Y. R. et al. Self-assembling human heart organoids for the modeling of cardiac development and congenital heart disease. Nat Commun 12, 5142 (2021).

34. Hofbauer, P. et al. Cardioids reveal self-organizing principles of human cardiogenesis. Cell 184, 3299–3317.e22 (2021).

35. Rood, J. E. et al. The Human Cell Atlas from a cell census to a unified foundation model. Nature 637, 1065–1071 (2025).

36. Knight-Schrijver, V. et al. A human developing heart atlas reveals cardiovascular cell diversity for in vitro model benchmarking. Preprint at 10.1101/2024.04.27.591127 (2024).

37. Organoid Network - Overview - HCA Data Portal. https://data.humancellatlas.org/hca-bio-networks/organoid.

38. He, Z. et al. An integrated transcriptomic cell atlas of human neural organoids. Nature 635, 690–698 (2024).

39. Xu, Q. et al. An integrated transcriptomic cell atlas of human endoderm-derived organoids. Nat Genet 57, 1201–1212 (2025).

40. Mueller, M. F. et al. Building optimized single-cell reference atlases with scAtlasTb. 2026.07.30.741695 Preprint at 10.64898/2026.07.30.741695 (2026)

41. Luecken, M. D. et al. Benchmarking atlas-level data integration in single-cell genomics. Nat Methods 19, 41–50 (2022).

42. Karbassi, E. et al. Cardiomyocyte maturation: advances in knowledge and implications for regenerative medicine. Nat Rev Cardiol 17, 341–359 (2020).

43. Browaeys, R. et al. MultiNicheNet: a flexible framework for differential cell-cell communication analysis from multi-sample multi-condition single-cell transcriptomics data. Preprint at 10.1101/2023.06.13.544751 (2023).

44. Sun, J. et al. hapln1 Defines an Epicardial Cell Subpopulation Required for Cardiomyocyte Expansion During Heart Morphogenesis and Regeneration. Circulation 146, 48–63 (2022).

45. Mazzotta, S. et al. Distinctive Roles of Canonical and Noncanonical Wnt Signaling in Human Embryonic Cardiomyocyte Development. Stem Cell Reports 7, 764–776 (2016).

46. Bhattacharya, A. et al. Myocardial TGFβ2 Is Required for Atrioventricular Cushion Remodeling and Myocardial Development. Journal of Cardiovascular Development and Disease 8, 26 (2021).

47. Dogra, D. et al. Opposite effects of Activin type 2 receptor ligands on cardiomyocyte proliferation during development and repair. Nat Commun 8, 1902 (2017).

48. Chanthra, N. et al. A Novel Fluorescent Reporter System Identifies Laminin-511/521 as Potent Regulators of Cardiomyocyte Maturation. Sci Rep 10, 4249 (2020).

49. Pollitt, E. J. G., Sánchez-Posada, J., Snashall, C. M., Derrick, C. J. & Noël, E. S. Llgl1 mediates timely epicardial emergence and establishment of an apical laminin sheath around the trabeculating cardiac ventricle. Development 151, dev202482 (2024).

50. Ton, M.-L. N. et al. An atlas of rabbit development as a model for single-cell comparative genomics. Nat Cell Biol 25, 1061–1072 (2023).

51. Andreatta, M. & Carmona, S. J. UCell and pyUCell: single-cell gene signature scoring for R and Python. Bioinformatics 42, btag055 (2026).

52. Hrovatin, K. et al. Integrating single-cell RNA-seq datasets with substantial batch effects. BMC Genomics 26, 974 (2025).

53. Kubalak, S. W., Miller-Hance, W. C., O’Brien, T. X., Dyson, E. & Chien, K. R. Chamber specification of atrial myosin light chain-2 expression precedes septation during murine cardiogenesis. J Biol Chem 269, 16961–16970 (1994).

54. O’Brien, T. X., Lee, K. J. & Chien, K. R. Positional specification of ventricular myosin light chain 2 expression in the primitive murine heart tube. Proc Natl Acad Sci U S A 90, 5157–5161 (1993).

55. Argelaguet, R. et al. Multi-Omics Factor Analysis-a framework for unsupervised integration of multi-omics data sets. Mol Syst Biol 14, e8124 (2018).

56. Dann, E., Henderson, N. C., Teichmann, S. A., Morgan, M. D. & Marioni, J. C. Differential abundance testing on single-cell data using k-nearest neighbor graphs. Nat Biotechnol 40, 245–253 (2022).

57. Perl, E. & Waxman, J. S. Reiterative Mechanisms of Retinoic Acid Signaling during Vertebrate Heart Development. J Dev Biol 7, 11 (2019).

58. Li, J., Yue, Y. & Zhao, Q. Retinoic Acid Signaling Is Essential for Valvulogenesis by Affecting Endocardial Cushions Formation in Zebrafish Embryos. Zebrafish 13, 9–18 (2016).

59. Zou, X. et al. Self-assembled chamber-like cardiac organoids for modeling cardiac chamber formation and cardiotoxicity assessment. Nat Commun 17, 7096 (2026).

60. Lotfollahi, M. et al. Mapping single-cell data to reference atlases by transfer learning. Nat Biotechnol 40, 121–130 (2022).

61. Domínguez Conde, C., et al. Cross-tissue immune cell analysis reveals tissue-specific features in humans. Science 376, eabl5197 (2022).

62. Sikkema, L. et al. Automated evaluation of single-cell reference atlas mappings enables the identification of disease-associated cell states.b Preprint at 10.1101/2025.05.23.655749 (2025).

63. CZI Single-Cell Biology et al. CELLxGENE: a performant, scalable exploration platform for high dimensional sparse matrices. Preprint at 10.1101/2021.04.05.438318 (2021).

64. Lotfollahi, M. et al. ArchMap is a web-based platform for reference-based analysis of single-cell datasets. Nat Genet 58, 2469–2473 (2026).

65. Richards, D. J. et al. Human cardiac organoids for the modelling of myocardial infarction and drug cardiotoxicity. Nat Biomed Eng 4, 446–462 (2020).

66. Chamberlin, J. T., Lee, Y., Marth, G. T. & Quinlan, A. R. Differences in molecular sampling and data processing explain variation among single-cell and single-nucleus RNA-seq experiments. Genome Research 34, 179–188 (2024).

67. Tirgar, P. et al. Heart-on-a-Miniscope: A Miniaturized Solution for Electrophysiological Drug Screening in Cardiac Organoids. Small 21, 2409571 (2025).

68. Ernst, P., Bidwell, P. A., Dora, M., Thomas, D. D. & Kamdar, F. Cardiac calcium regulation in human induced pluripotent stem cell cardiomyocytes: Implications for disease modeling and maturation. Front. Cell Dev. Biol. 10, (2023).

69. Hou, C. et al. iPSC-derived cardiac organoids for drug cardiotoxicity evaluation and efficacy prediction in myocardial infarction and cardiac hypertrophy models. Stem Cell Res Ther 17, 296 (2026).

70. Ahmed, R. E., Anzai, T., Chanthra, N. & Uosaki, H. A Brief Review of Current Maturation Methods for Human Induced Pluripotent Stem Cells-Derived Cardiomyocytes. Front. Cell Dev. Biol. 8, (2020).

71. Weinberger, M., Simões, F. C., Patient, R., Sauka-Spengler, T. & Riley, P. R. Functional Heterogeneity within the Developing Zebrafish Epicardium. Developmental Cell 52, 574–590.e6 (2020).

72. Braitsch, C. M., Combs, M. D., Quaggin, S. E. & Yutzey, K. E. Pod1/Tcf21 is regulated by retinoic acid signaling and inhibits differentiation of epicardium-derived cells into smooth muscle in the developing heart. Developmental Biology 368, 345–357 (2012).

73. Lupu, I.-E., Redpath, A. N. & Smart, N. Spatiotemporal Analysis Reveals Overlap of Key Proepicardial Markers in the Developing Murine Heart. Stem Cell Reports 14, 770–787 (2020).

74. Gambardella, L. et al. BNC1 regulates cell heterogeneity in human pluripotent stem cell-derived epicardium. Development 146, dev174441 (2019).

75. Lie-Venema, H. et al. Origin, fate, and function of epicardium-derived cells (EPDCs) in normal and abnormal cardiac development. ScientificWorldJournal 7, 1777–1798 (2007).

76. Acharya, A. et al. The bHLH transcription factor Tcf21 is required for lineage-specific EMT of cardiac fibroblast progenitors. Development 139, 2139–2149 (2012).

77. Su, T. et al. Single-cell analysis of early progenitor cells that build coronary arteries. Nature 559, 356–362 (2018).

78. Quintard, C. et al. A microfluidic platform integrating functional vascularized organoids-on-chip. Nat Commun 15, 1452 (2024).

79. Wiesinger, A. et al. A single cell transcriptional roadmap of human pacemaker cell differentiation. eLife 11, e76781 (2022).

80. Iyer, D. et al. Robust derivation of epicardium and its differentiated smooth muscle cell progeny from human pluripotent stem cells. Development 142, 1528–1541 (2015).

81. Protze, S. I. et al. Sinoatrial node cardiomyocytes derived from human pluripotent cells function as a biological pacemaker. Nat Biotechnol 35, 56–68 (2017).

82. Ginistrelli, L. C. et al. Dissecting human fetal cardiac repair using cardioids. Preprint at 10.64898/2026.06.30.735236 (2026).

83. Fischer, B. et al. A complete workflow for the differentiation and the dissociation of hiPSC-derived cardiospheres. Stem Cell Research 32, 65–72 (2018).

84. Wolf, F. A., Angerer, P. & Theis, F. J. SCANPY: large-scale single-cell gene expression data analysis. Genome Biol 19, 15 (2018).

85. Wolock, S. L., Lopez, R. & Klein, A. M. Scrublet: Computational Identification of Cell Doublets in Single-Cell Transcriptomic Data. Cell Systems 8, 281–291.e9 (2019).

86. Lopez, R., Regier, J., Cole, M. B., Jordan, M. I. & Yosef, N. Deep generative modeling for single-cell transcriptomics. Nat Methods 15, 1053–1058 (2018).

87. Xu, C. et al. Probabilistic harmonization and annotation of single-cell transcriptomics data with deep generative models. Mol Syst Biol 17, e9620 (2021).

88. De Donno, C. et al. Population-level integration of single-cell datasets enables multi-scale analysis across samples. Nat Methods 20, 1683–1692 (2023).

89. Korsunsky, I. et al. Fast, sensitive and accurate integration of single-cell data with Harmony. Nat Methods 16, 1289–1296 (2019).

90. Palarea-Albaladejo, J. & Martín-Fernández, J. A. zCompositions — R package for multivariate imputation of left-censored data under a compositional approach. Chemometrics and Intelligent Laboratory Systems 143, 85–96 (2015).

91. van den Boogaart, K. G. & Tolosana-Delgado, R. “compositions”: A unified R package to analyze compositional data. Computers & Geosciences 34, 320–338 (2008).

92. Robinson, M. D., McCarthy, D. J. & Smyth, G. K. edgeR: a Bioconductor package for differential expression analysis of digital gene expression data. Bioinformatics 26, 139–140 (2010).

93. Robinson, M. D. & Oshlack, A. A scaling normalization method for differential expression analysis of RNA-seq data. Genome Biol 11, R25 (2010).

94. Lund, S. P., Nettleton, D., McCarthy, D. J. & Smyth, G. K. Detecting differential expression in RNA-sequence data using quasi-likelihood with shrunken dispersion estimates. Stat Appl Genet Mol Biol 11, /j/sagmb.2012.11.issue-5/1544-6115.1826/1544-6115.1826.xml (2012).

95. Fang, Z., Liu, X. & Peltz, G. GSEApy: a comprehensive package for performing gene set enrichment analysis in Python. Bioinformatics 39, btac757 (2023).

96. Amezquita, R. A. et al. Orchestrating single-cell analysis with Bioconductor. Nat Methods 17, 137–145 (2020).

97. Browaeys, R., Saelens, W. & Saeys, Y. NicheNet: modeling intercellular communication by linking ligands to target genes. Nat Methods 17, 159–162 (2020).

98. Murphy, S. A., Chen, E. Z., Tung, L., Boheler, K. R. & Kwon, C. Maturing heart muscle cells: Mechanisms and transcriptomic insights. Semin Cell Dev Biol 119, 49–60 (2021).

99. Giacomelli, E. et al. Human-iPSC-Derived Cardiac Stromal Cells Enhance Maturation in 3D Cardiac Microtissues and Reveal Non-cardiomyocyte Contributions to Heart Disease. Cell Stem Cell 26, 862–879.e11 (2020).

100. Eldemire, R., Tharp, C. A., Taylor, M. R. G., Sbaizero, O. & Mestroni, L. The Sarcomeric Spring Protein Titin: Biophysical Properties, Molecular Mechanisms, and Genetic Mutations Associated with Heart Failure and Cardiomyopathy. Curr Cardiol Rep 23, 121 (2021).

101. England, J. et al. Tropomyosin 1: Multiple roles in the developing heart and in the formation of congenital heart defects. Journal of Molecular and Cellular Cardiology 106, 1–13 (2017).

102. Miklas, J. W. et al. TFPa/HADHA is required for fatty acid beta-oxidation and cardiolipin re-modeling in human cardiomyocytes. Nat Commun 10, 4671 (2019).

103. Dufour, C. R. et al. Genome-wide Orchestration of Cardiac Functions by the Orphan Nuclear Receptors ERRα and γ. Cell Metabolism 5, 345–356 (2007).

104. Guo, Y. & Pu, W. T. Cardiomyocyte Maturation: New Phase in Development. Circulation Research 126, 1086–1106 (2020).

105. Gayoso, A. et al. A Python library for probabilistic analysis of single-cell omics data. Nat Biotechnol 40, 163–166 (2022).

106. Heumos, L. et al. Pertpy: an end-to-end framework for perturbation analysis. Nat Methods 23, 350–359 (2026).

107. Yu, G., Wang, L.-G., Han, Y. & He, Q.-Y. clusterProfiler: an R Package for Comparing Biological Themes Among Gene Clusters. OMICS 16, 284–287 (2012).

