## Supplementary material for "A Single-Cell Transcriptomic Atlas of Human Heart Organoids across Differentiation Protocols": Figure S1

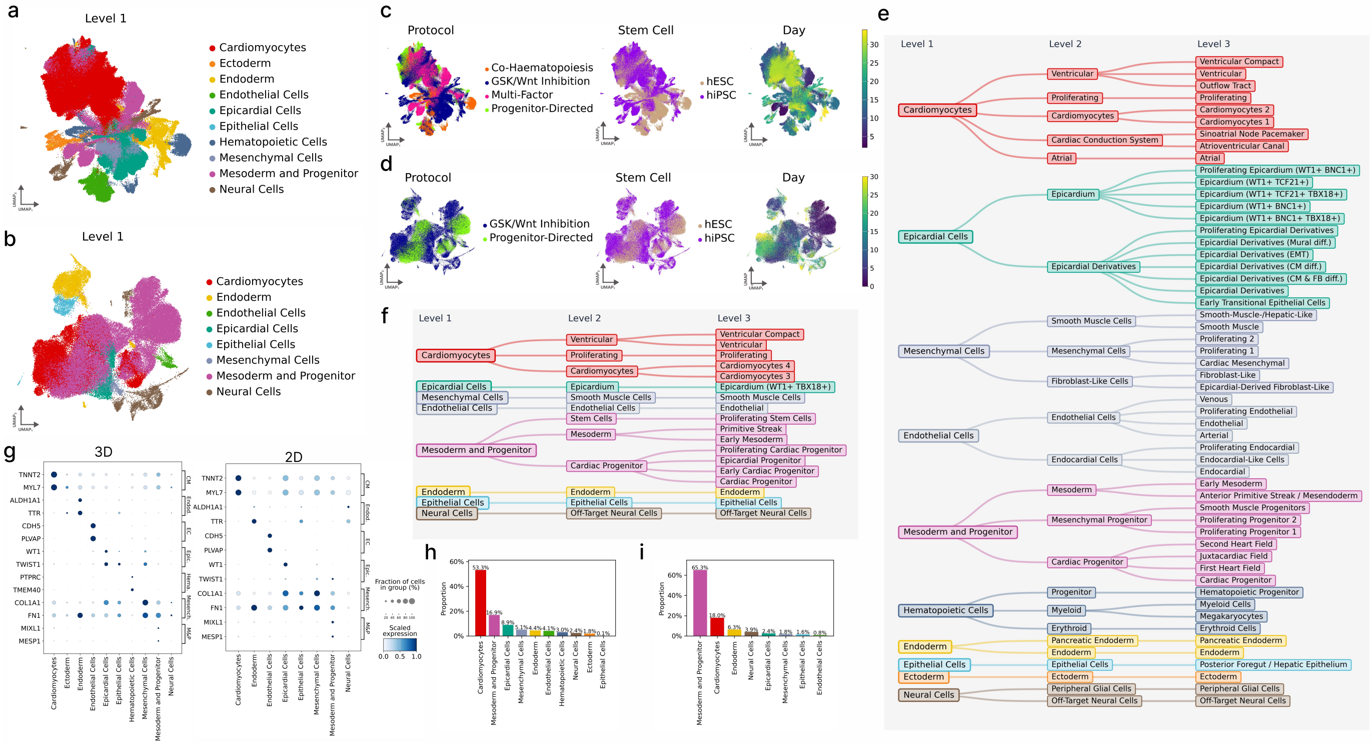


**Figure S1: 3D and 2D HHOCA atlases reveal a shared, multi-level cell type landscape of heart development in vitro**

**a,** UMAP embedding of the 3D organoid atlas, colored by level 1 cell type. **b,** UMAP embedding of the 2D monolayer atlas, colored by level 1 cell type. **c,** UMAP embeddings of the 3D atlas colored by protocol type, stem cell type, and day. **d,** UMAP embeddings of the 2D atlas colored by protocol type, stem cell type, and day. **e,** Hierarchical cell type annotation structure for the 3D atlas. **f,** Hierarchical cell type annotation structure for the 2D atlas. **g,** Dot plot of marker gene expression across organoids (left) and monolayers (right). **h,** Proportion of level 1 cell types in the 3D atlas. **i,** Proportion of level 1 cell types in the 2D atlas.
