## Supplementary material for "A Single-Cell Transcriptomic Atlas of Human Heart Organoids across Differentiation Protocols": Figure S2

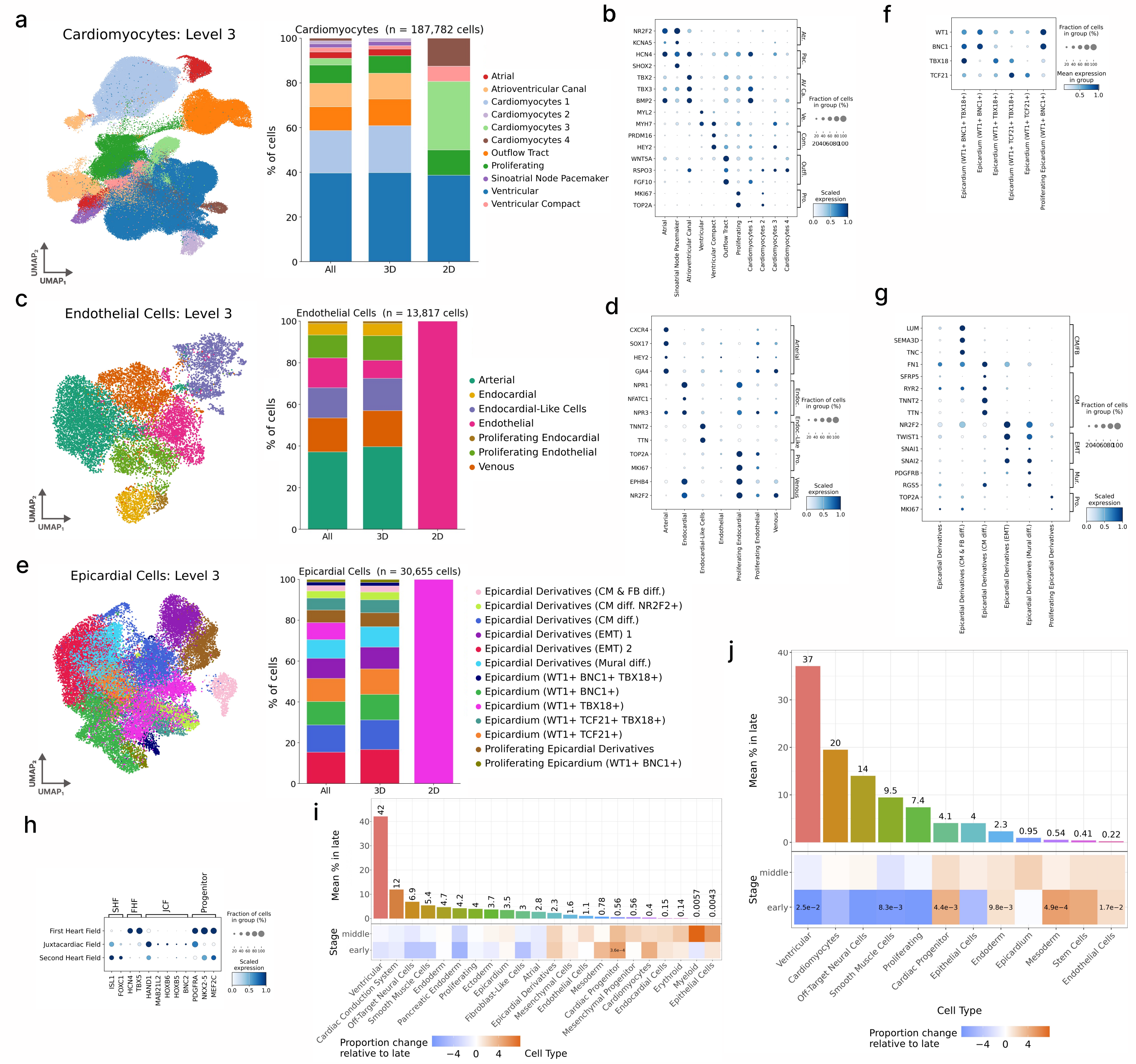

**Figure S2: Level 3 annotation reveals subtype-specific marker expression and composition differences between 3D and 2D**

**a,** UMAP embedding of cardiomyocytes colored by level 3 annotation (left), and proportion of level 3 cardiomyocyte subtypes across all, 3D, and 2D cells (right). **b,** Dot plot of marker gene expression across cardiomyocyte level 3 subtypes. **c,** UMAP embedding of endothelial cells colored by level 3 annotation (left), and proportion of level 3 endothelial subtypes across all, 3D, and 2D cells (right). **d,** Dot plot of marker gene expression across endothelial level 3 subtypes. **e,** UMAP embedding of epicardial cells colored by level 3 annotation (left), and proportion of level 3 epicardial subtypes across all, 3D, and 2D cells (right). **f,** Dot plot of epicardial specification marker expression (WT1, BNC1, TBX18, TCF21) across epicardial subtypes. **g,** Dot plot of marker gene expression across epicardial derivative subtypes. **h,** Dot plot of first and second heart field (FHF/SHF) and progenitor marker gene expression, showing distinct expression profiles between heart field progenitor populations. **i,** Changes in cell-type composition (cell_type_level2) across developmental stages in 3D cultures. Top: mean percentage of each cell type in the late stage. Bottom: heatmap of the log2 fold changes in mean cell-type proportion in the early and middle stages relative to late (orange = higher, blue = lower). Tile numbers are nominal p-values from linear models on CLR-transformed proportions, shown where the Benjamini-Hochberg FDR is below 0.05. Cell types are ordered by their abundance in the late stage. **j,** Same analysis as in (i), for 2D cultures.
