## Supplementary material for "A Single-Cell Transcriptomic Atlas of Human Heart Organoids across Differentiation Protocols": Figure S3

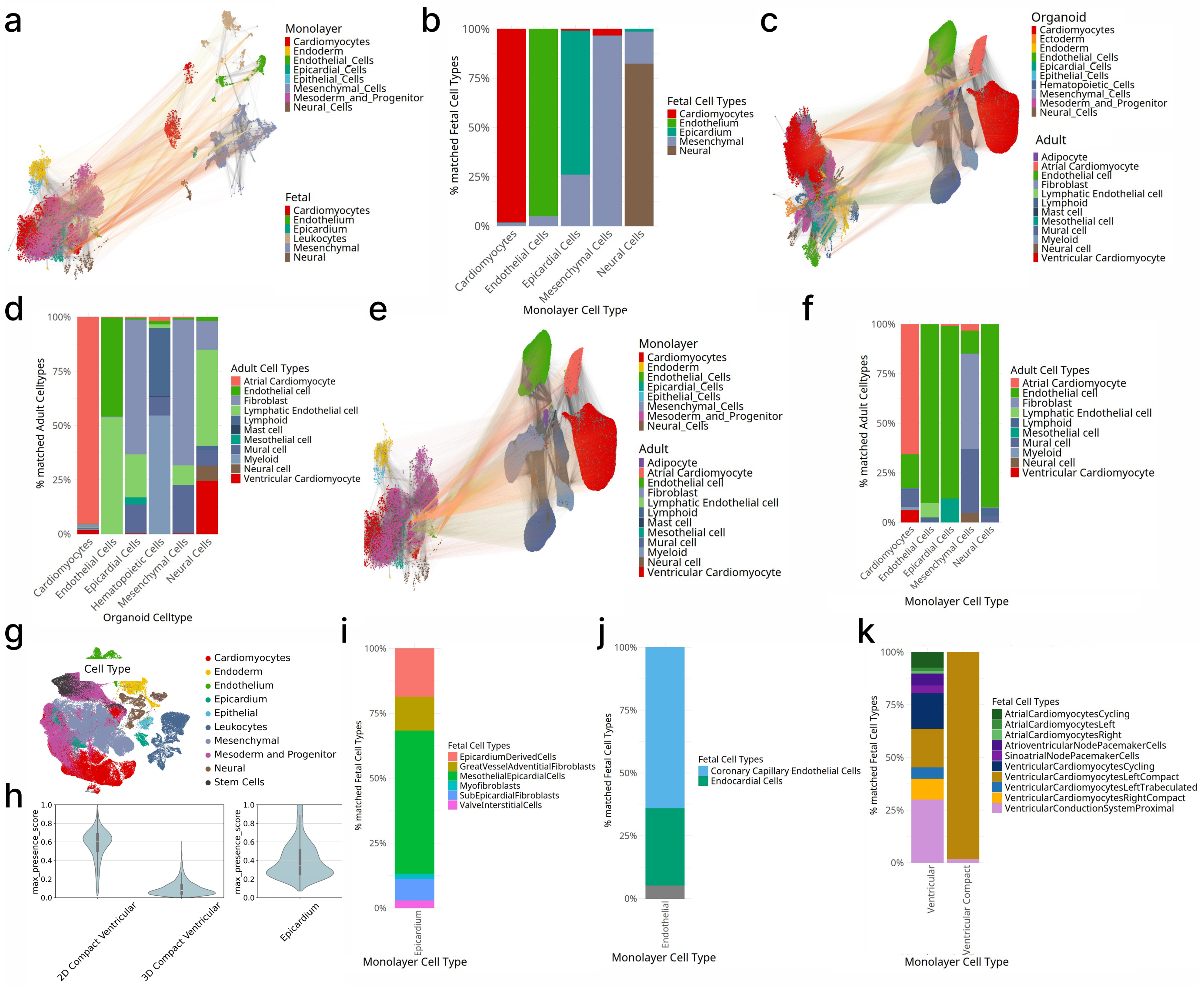


**Figure S3: Monolayers resemble fetal heart tissue, yet adult heart correspondence remains limited for monolayers and organoids.**

**a,** Neighborhood matching between the monolayer atlas and the fetal heart atlas **b,** Proportion of matched fetal cell types for each monolayer cell type, showing the correspondence between broad monolayer and fetal cell type annotations. **c,** Neighborhood matching between the organoid atlas and the adult heart atlas. **d,** Proportion of matched adult cell types for each organoid cell type. **e,** Neighborhood matching between the monolayer atlas and the adult heart atlas. **f,** Proportion of matched adult cell types for each monolayer cell type. **g,** Integrated embedding of monolayer and fetal cells, colored by cell type. **h,** Violin plot with boxplot of presence scores of compact cardiomyocytes and of monolayer epicardial cells. **i,** Proportion of matched fetal epicardial subtypes for monolayer epicardium. **j,** Proportion of matched fetal endothelial/endocardial subtypes for monolayer endothelium. **k,** Proportion of matched fetal cardiomyocyte subtypes for monolayer ventricular and ventricular compact cardiomyocytes.
