## Supplementary material for "A Single-Cell Transcriptomic Atlas of Human Heart Organoids across Differentiation Protocols": Figure S4

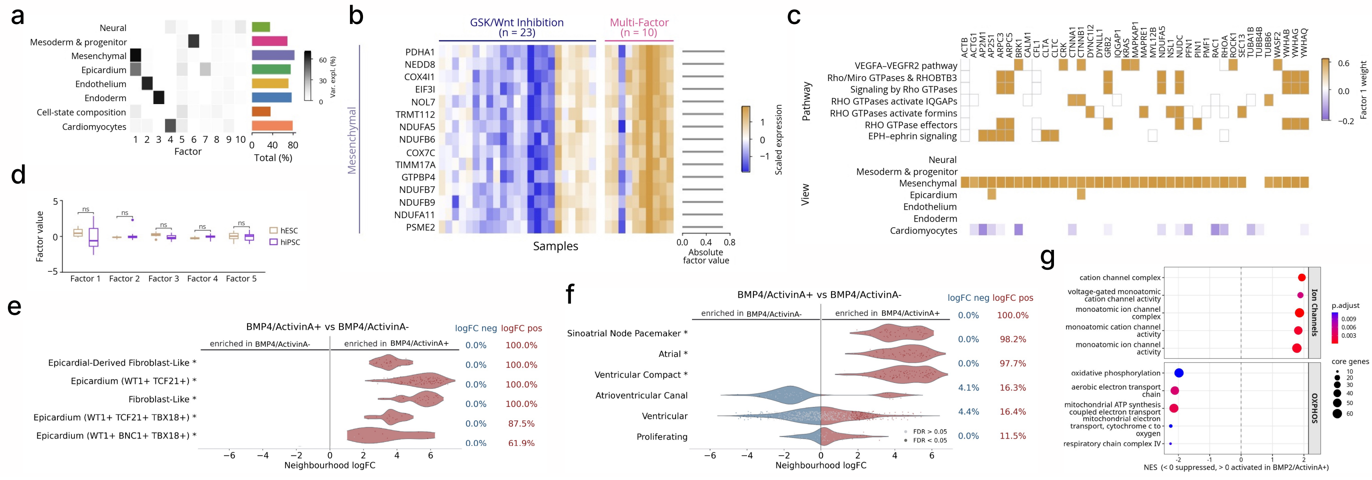


**Figure S4: MOFA and differential abundance analysis reveals protocol-associated gene programs and BMP4/Activin A-driven cell type shifts**

**a,** Variance explained by each MOFA factor across cell type views with total explained variance per factor shown on the right. **b,** Heatmap of normalized expression values for the top 15 features (mesenchymal view) contributing to Factor 1, shown across samples grouped by protocol (GiWi vs. Multi-Factor), with absolute factor loadings shown on the right. **c,** Heatmap of pathway enrichment among the top Factor 1 genes across cell type views, showing enriched pathways (top) and their contributing genes per view (bottom), with the Mesenchymal view driving the strongest associations. **d,** Boxplots comparing MOFA factor values (Factors 1-5) between hESC- and hiPSC-derived sample, with significance indicated above each comparison (ns = not significant) **e,** Differential abundance of epicardial and fibroblast-related subtypes and **f,** cardiomyocyte subtypes between BMP4/Activin A+ and BMP4/Activin A- organoids, tested using Milo. Percentages of significant neighborhoods (FDR < 0.05) enriched in each direction are shown on the right. **g,** Gene set enrichment analysis of genes activated or suppressed in BMP4/Activin A+ relative to BMP4/Activin A- GiWi protocols, grouped by biological theme.
