## Supplementary material for "A Single-Cell Transcriptomic Atlas of Human Heart Organoids across Differentiation Protocols": Figure S5

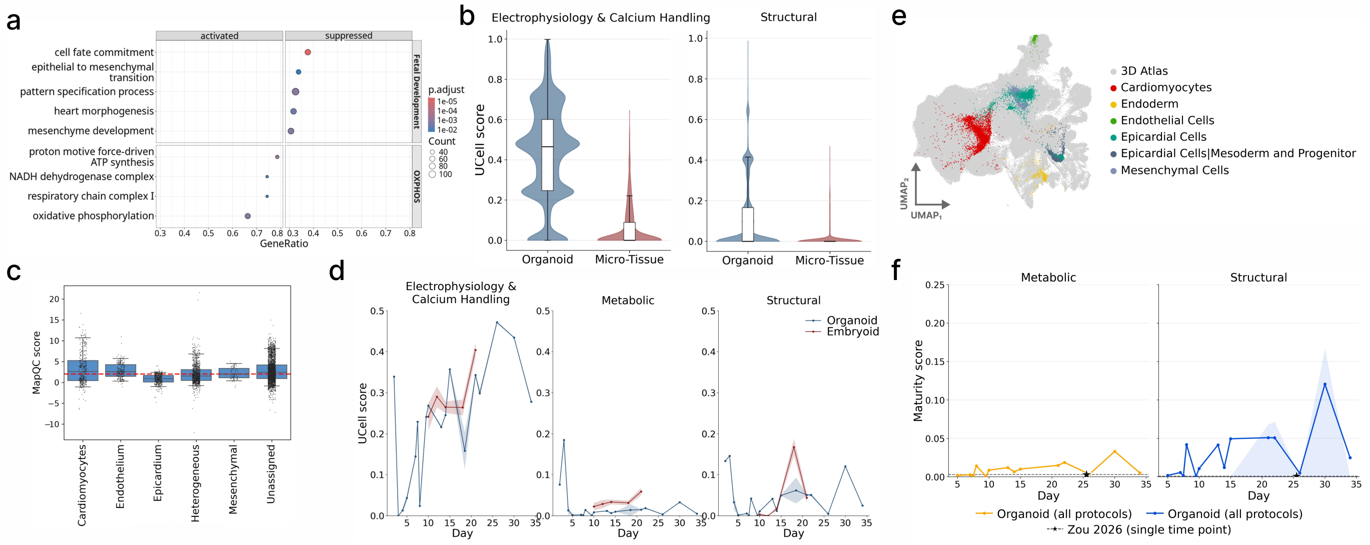


### Figure S5: Atlas-based benchmarking reveals maturity differences across diverse cardiac in vitro systems

**a,** Gene set enrichment analysis of pathways activated or suppressed in micro-tissue mesenchymal cells relative to organoids. **b,** UCell scores for electrophysiology/calcium handling and structural maturity gene sets in organoid day 30 versus micro-tissue day 34 cardiomyocytes. **c,** MapQC scores for embryoid body cell types mapped onto the organoid atlas, grouped by majority-voted cell type; dashed line indicates the quality threshold (MapQC score = 2). **d,** UCell scores for electrophysiology/calcium handling, metabolic, and structural maturity gene sets across organoid and embryoid body cardiomyocytes over time (day). **e,** UMAP embedding of the atlas (gray) with the query organoid dataset (Zou et al.) mapped on, colored by predicted cell type. **f,** Metabolic and structural maturity scores across organoid day, comparing all organoid protocols in the atlas (blue line) to the query organoid dataset (Zou et al., dashed line/black star).
