## Supplementary material for "A Single-Cell Transcriptomic Atlas of Human Heart Organoids across Differentiation Protocols": Figure S6

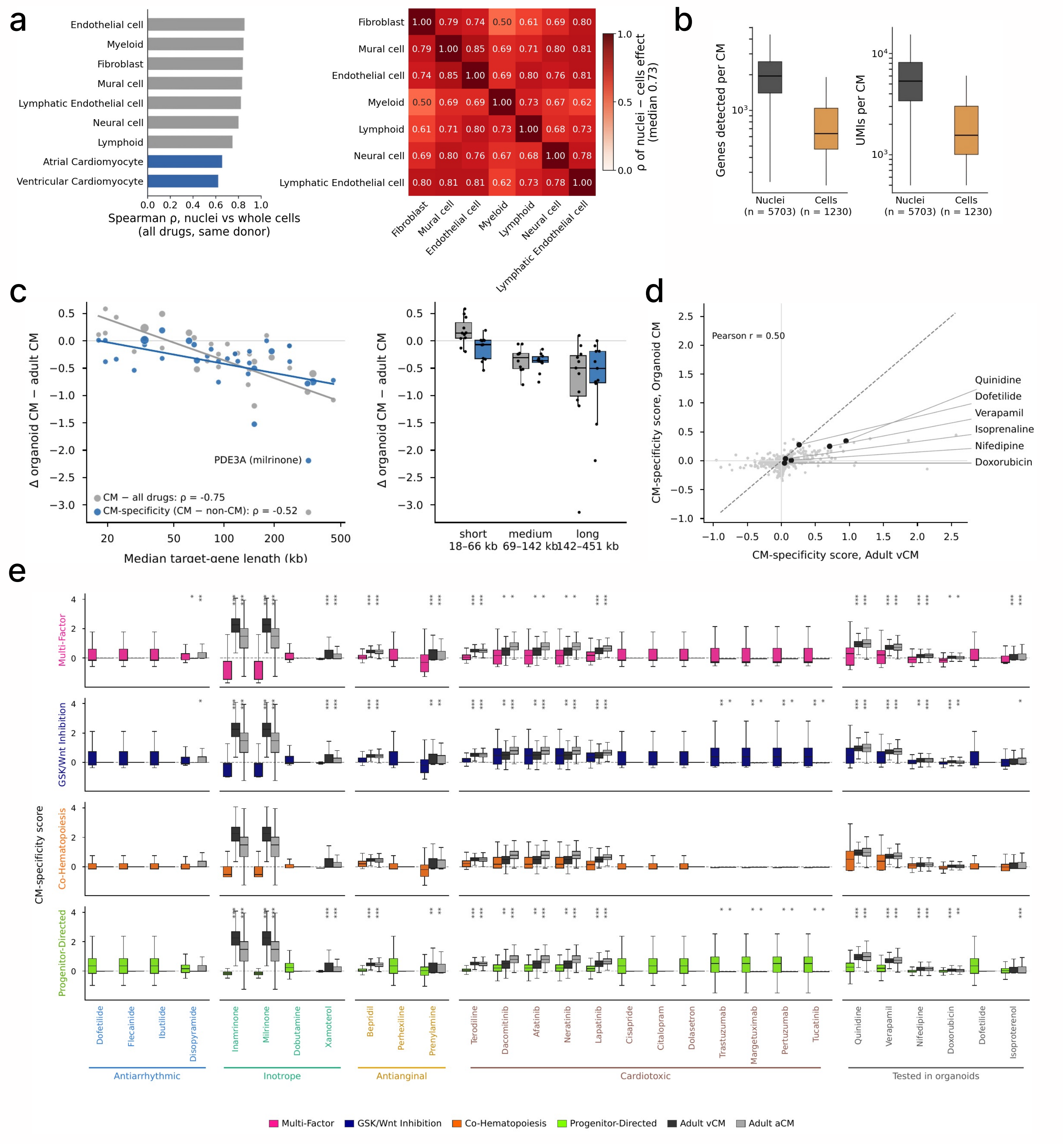


**Figure S6: Reduced organoid drug-target specificity correlates with transcript length but is not fully explained by nuclei-versus-whole-cell sequencing bias**

**a,** Left: Spearman correlation of drug scores between matched nuclei and cells, per cell type. Right: correlation of the nuclei − cells difference across cell types (median ρ = 0.73). **b,** Genes and UMIs per CM in nuclei (n = 5,703) vs cells (n = 1,230). **c,** The organoid − adult CM score difference (Δ) per drug becomes more negative as median target-gene length increases (Spearman ρ = −0.75 for CM score, −0.52 for CM-specificity). Right: drugs grouped into target-length bins. **d,** CM-specificity scores in adult vCM vs organoid CM for all drugs (Pearson r = 0.50). The dashed line marks y = x. Drugs previously tested in organoids are labelled. **e,** CM-specificity scores (CM minus mean non-CM score) for drugs by cardiac class, and for drugs previously tested in organoids. Rows show organoid differentiation protocols; colored boxes are organoid CMs, grey boxes are adult vCM and aCM. Asterisks show organoid vs vCM (left) and vs aCM (right), two-sided Mann–Whitney U test with Benjamini–Hochberg correction. *q < 0.05, **q < 0.01, ***q < 0.001; ns, not significant; n/a, not tested.
